# Vimentin bridges scales to convert polarized cell locomotion into coordinated collective migration

**DOI:** 10.64898/2026.01.12.699001

**Authors:** Sudheer Kumar Peneti, Satish Babu Morparthi, Clémence Thiant, Joseph d’Alessandro, Lucas Anger, Ranjith Chilupuri, Tien Dang, Manon Arnaud, Julia Eckert, Stéphane Vassilopoulos, Benoit Ladoux, René-Marc Mège

**Author notes:** Correspondance &.

## Abstract

Collective cell migration is central in development and disease. Vimentin is an intermediate filament protein expressed by epithelial cells at the edge of wounds where collective cell migration is most efficient. Yet, its functional role in this context remains underexplored. Here, we show that vimentin, over-expressed in cells undergoing partial epithelial to mesenchymal transition at the edge of epithelial monolayers, has a multiscale impact on the whole monolayer mechano-dynamics. Vimentin knock-down delays wound closure, reduces cell coordination, while increasing traction forces exerted by cells on the substratum. It also disrupts the directionality of leader cells migration, as well as the cohesion and coordinated motion of cells deep in the monolayer. We further show that vimentin promotes the conversion of polarized cell locomotion into coordinate collective migration by polarizing actin, focal adhesions and traction forces, sustaining leader cell’s lamellipodium protrusive activity and directionality, while allowing mechanical coupling of leader with follower cells. Altogether, we show that vimentin is essential for bridging polarized single cell locomotion and coordinated collective migration to allow efficient collective migration.

**Significant statement:** Epithelial cells migrate coordinately to repair tissue. Vimentin, transiently enriched at the wound edge plays a key role in guiding this process by converting polarized cell locomotion into coordinate collective migration, enabling leader cells to polarize, form stable protrusions, and maintain directed migration, while allowing mechanical coupling of front cells with followers, necessary for efficient coordinated collective migration. These findings reveal vimentin, a known EMT marker, as a novel positive regulator of efficient wound healing.

## INTRODUCTION

Collective cell migration is a fundamental process in embryogenesis, metastasis, and tissue repair. At the single cell level, migration is achieved by symmetry breaking and acquiring a front-rear polarity with the formation of a protruding lamellipodial structure at the front and a retracting edge at the rear (Pankov et al., 2005; Petrie et al., 2009; Ridley et al., 2003). This process involves the coordination between actomyosin-generated traction forces, activities of integrin-mediated cell-matrix adhesions that provide friction to the substratum, and actin polymerization ahead of adhesions that promotes forward protrusion (Ladoux et al., 2016; Petrie et al., 2009; Ridley et al., 2003). Mechanisms that generate tractions exhibit a ‘force dipole’ that propels the cell in a forward direction (Ladoux & Mège, 2017). This cell-autonomous propulsion is observed across a wide range of mesenchymal and epithelial cell types (De Rooij et al., 2005; Pankov et al., 2005; Plotnikov et al., 2012). However, within epithelial monolayers, cells are additionally interconnected through cell–cell junctions that couple the cells mechanically allowing tension building in the monolayer, and allow cells to coordinate their migration over large scales (Capuana et al., 2020; Mayor & Etienne-Manneville, 2016; Petitjean et al., 2010; Trepat et al., 2009). During wound healing in MDCK monolayers, dynamic instabilities at the leading edge give rise to leader cells that further organize multicellular structures called “fingers” (Poujade et al., 2007; Reffay et al., 2014; Vishwakarma et al., 2018). Leader cells at the fingertip have distinct morphological and mechanical properties with a large lamellipodium, high protrusive activity and the highest traction forces. These cells are essential not only for the formation of the finger, but also to transmit mechanical forces to their followers, guiding them such as finger forms a mechanical unit in which cells migrate together as a single global entity with coordinated directionality and polarity (Khalil & Friedl, 2010; Reffay et al., 2011, 2014). Mechanical cues and biochemical signals are integrated to coordinate tissue-scale polarization in the migrating monolayers as well as in fingers (Reffay et al., 2014; Sunyer et al., 2016; Vedula et al., 2012). So far, only a very limited number of mechanochemical pathways have been reported to be essential for collective epithelial cell migration. These include pathways associated with cell-cell adhesion complexes and the actomyosin cytoskeleton (Bazellières et al., 2015; Ladoux et al., 2010, 2010) as well as the propagation of extracellular signal-regulated kinase (ERK) activation waves reported to yield long-distance transmission of guidance cues regulating collective cell polarization and migration (Aoki et al., 2017; Hino et al., 2020; Matsubayashi et al., 2004).

Vimentin is a type III intermediate filament protein, prominently expressed in mesenchymal cells (Gruenbaum & Aebi, 2014; D.-H. Kim & Wirtz, 2013; J. Kim et al., 2016), where it plays a critical role in single cell mechanics and potentially directed migration (Costigliola et al., 2017; Gan et al., 2016). Alteration in the vimentin network in single cells has been shown to influence actomyosin and microtubule networks (Jiu et al., 2015, 2017) (Alisafaei et al., 2024), focal adhesion dynamics (Burgstaller et al., 2010; Eckes et al., 1998; Gregor et al., 2014; Tsuruta & Jones, 2003). In astrocytes, the silencing of all three type III filaments (vimentin, nestin, and GFAP) affects their collective cell migration (De Pascalis et al., 2018). Vimentin is a well-recognized Epithelial to Mesenchymal Transition (EMT) and invasive cancer cells biomarker (Satelli & Li, 2011). During epithelial wound healing, edge cells which undergo partial EMT and upregulate vimentin expression (Gayrard et al., 2018; Gilles et al., 1999; Menko et al., 2014). In MCF10A mammary epithelial cells, increased expression of vimentin at the leading edge has been associated with increased migration speed and wound healing efficiency (Gilles et al., 1999). However, the precise role of vimentin in controlling the dynamics and mechanics of epithelial collective cell migration remains poorly understood.

Here, we addressed the role of vimentin in the dynamics and mechanics of epithelial collective migration, using a well-controlled wound healing assay in MDCK monolayers. We show that vimentin knockout slows down wound healing, increases cell substratum adhesion and traction forces in the whole monolayer, and impairs coordinated, directed cell migration. Moreover, vimentin loss of function has a major impact on the formation, coordination and directed migration of fingers. In the absence of vimentin overexpression, we observe a disorganization of the actin arcs in leader cells, accompanied by a redistribution of traction forces and altered lamellipodia dynamics. Thus, vimentin expression is critical for maintaining leader cell protrusive activity and persistence, as well as for mechanical coupling and collective coordination of cells deep in the monolayer. Altogether, vimentin is essential for bridging polarized single cell locomotion and coordinated collective migration.

## RESULTS

### Vimentin expression increases at the leading edge of epithelial wounds

To address the role of vimentin in epithelial collective cell migration, we performed wound healing assays on MDCK cell monolayers **(Figure 1A)**. We examined the spatiotemporal expression pattern of vimentin. Immunostaining for vimentin, performed before the onset of collective migration (t=0 h, immediately after removal of the Polydimethylsiloxane [PDMS] block) and at t=24 h, revealed an increased expression of vimentin at the leading edge of the monolayer, which progressively decreased away from the edge **(Figure 1B and 1D; S1A)**. Conversely, the intensity of F-actin increased slightly from the leading edge to the bulk of the monolayer at t=24 h **(Figure 1B (lower panel) and 1D)**. In contrast, as expected for edge cells undergoing partial EMT, immunostaining revealed decreased levels of cytokeratins K8/18 at the edge (**Figure S1F, G**). To determine whether the accumulation of vimentin was due to an increase of protein expression, induced by the presence of the free edge, we compared the expression of vimentin in sparse versus confluent cultures, and found a 3-fold increase in vimentin protein content in sparse cells **(Figure S1B, C)**. While vimentin expression in epithelial cells has been associated with the initiation of cell migration at the edge of epithelial monolayers in other cell line models (Gilles et al., 1999; Menko et al., 2014), we observed an upregulation of vimentin even before removal of the physical constraint, when cells were still tightly packed **(Figure S1E)**, contrasting with the homogenous low level of expression in confluent monolayers lacking a free edge (**Figure S1D**). This suggested that the presence of a free edge is sufficient to promote vimentin upregulation at the edge of the monolayer before overt cell spreading and migration. Interestingly, vimentin was significantly expressed at higher levels in leader cells and fingers compared to interfinger regions **(Figure 1C, and 1E)**. Thus, vimentin upregulation is not uniform along the boundary, but is spatially reinforced in leader/finger regions.

**Figure 1:**
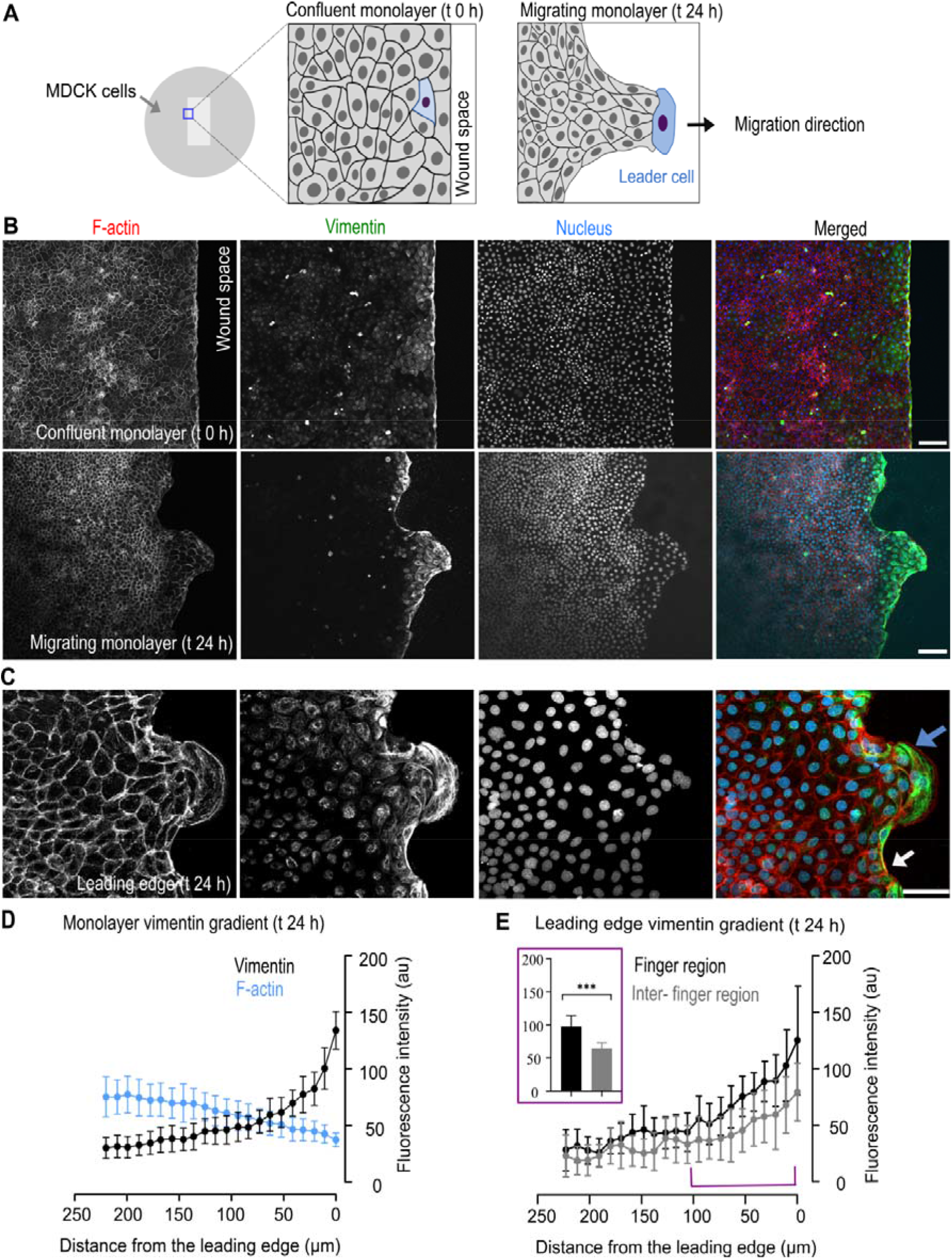
Upregulation of vimentin expression at the epithelial wound edge. **(A)** Schematic of the wound healing assay. MDCK cells were grown on the fibronectin-coated glass substrate overnight in the presence of a physical constraint up to reaching confluence. Then, the physical constraint was carefully removed, allowing cells to invade free space by collective cell migration, emergence of leader cells (indicated in blue) and fingers (t=24 h). **(B)** The images illustrate the distribution of actin, vimentin, and nuclei staining in the confluent monolayers prior to the onset of migration (t=0 h) and after 24 hours. Scale bars, 100 µm. **(C)** Higher magnification images at 24 h showing actin, nuclei and vimentin distribution profiles within the finger (denoted by a blue arrow) and interfinger regions (indicated by a white arrow). Scale bars, 50 µm. **(D)** Average intensity (arbitrary units) values ± SD of vimentin and F-actin stainings plotted as a function of the distance from the monolayer edge, at t 24h, binned for every 15 µm. **(E)** Average values ± SD of vimentin staining intensity in finger and interfinger regions, binned for every 15 µm, plotted as a function of distance from edge. The inset shows the average intensity of vimentin staining in the finger and interfinger regions from the edge to 100μm (marked by colored brackets). Mann-Whitney comparison, n = 12 from three independent experiments.

### Vimentin expression ensures fast, coherent, and directional collective cell migration

To elucidate the contribution of this vimentin upregulation to the dynamics and mechanics of healing monolayers, MDCK vimentin knock-out (KO) cells were generated using CRISPR-Cas9 technology **(Figure S2A)**. Using time-lapse phase contrast microscopy, we then captured live images of migrating KO and wild-type (WT) parental cells monolayers at 10-minute intervals for 24 h **(Figure 2A, video 1)**. Tracking the displacement of the monolayer front revealed that KO cells lagged behind WT cells. Quantitative analysis of the advancement of the healing front revealed an immediate and linear progression of WT cells following the removal of the physical constraint, whereas KO monolayers displayed a slower front progression during the first 5-6 h, followed by a gradual acceleration phase, to reach a linear progression after 10 h that remained consistently reduced compared to WT cells **(Figure 2A)**.

**Figure 2:**
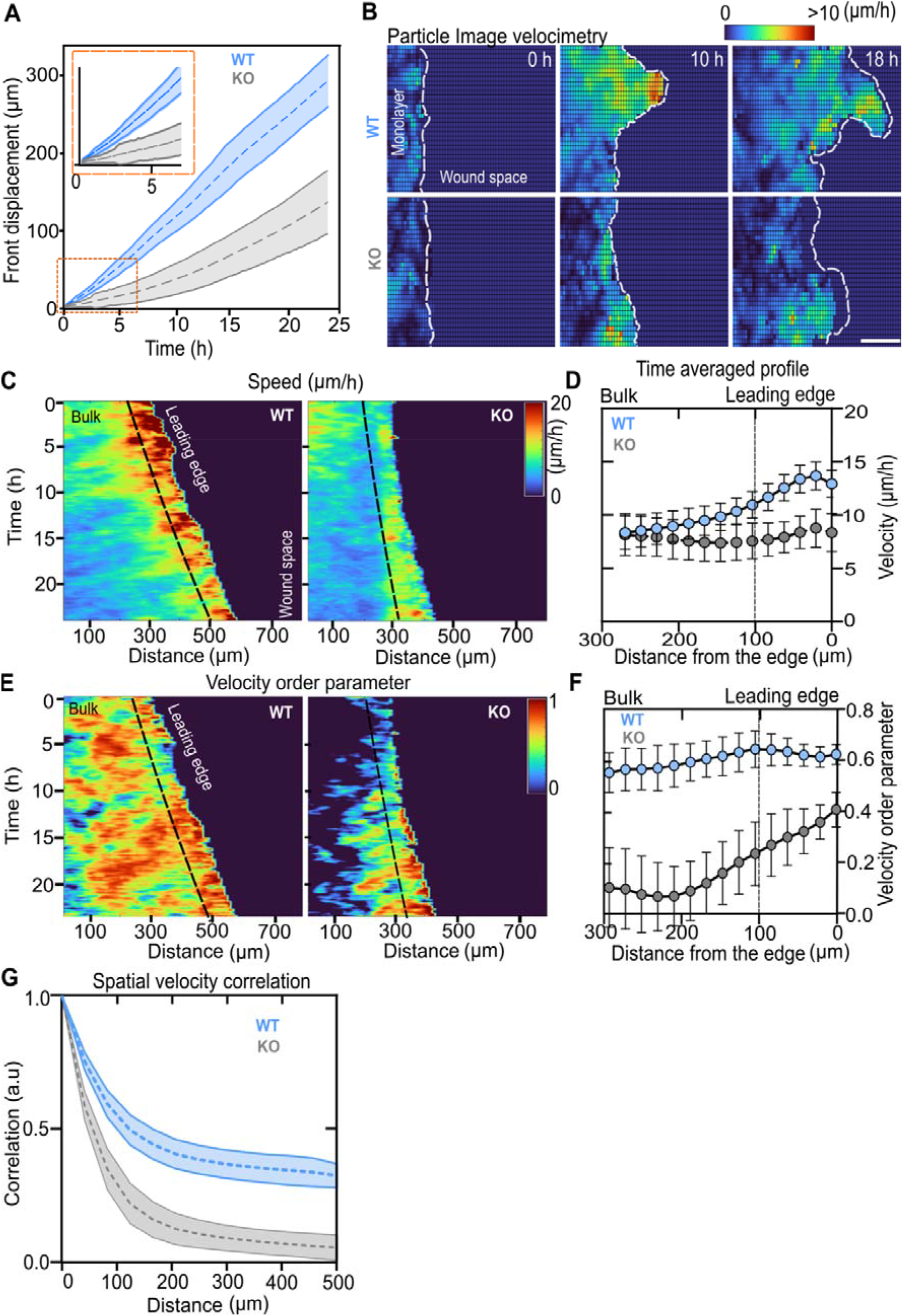
Loss of function of vimentin impairs coherent and directional collective cell migration. **(A)** Average displacement of the front migration over time. Insert (orange) represents initial displacements of monolayer from 0 to 5 hours. The dotted lines and shaded regions represent average and SDs. Sample sizes, n=13 monolayers (WT), n=16 monolayers (KO). **(B)** Velocity heat maps of the migrating monolayers over time. The images have been cropped to one-fourth of their original size for improved clarity. Scale bars, 100μm. **(C, E)** Kymographs representing speed (µm/hr) and order parameter over time for WT (left) and KO (right) monolayers. Red signifies high speed and strong alignment in the direction of migration, while blue indicates slower speeds and cell orientation opposing the direction of migration. **(D, F)** Time-averaged velocity and order parameter profiles as a function of the distance from the monolayer leading edge. Black thick line indicates the trend of averaged values. The filled blue and gray circles and black bars represent averages and SDs. The black dashed line indicates the position of the leading edge, separating the bulk of the monolayer from the wound space. Sample sizes n=13 monolayers (WT), and n=13 monolayers (KO). **(G)** Spatial velocity correlation function. The dotted lines indicate average values and shaded regions indicate SDs. Sample sizes n=13 monolayers (WT), n=16 monolayers (KO). Sample sizes were representative of three independent experiments.

Particle Image Velocimetry (PIV) was employed to generate velocity heatmaps for WT and KO monolayers **(Figure 2B, video 2, 3)**, from which kymographs of velocity profiles over time were extracted **(Figure 2C)**. The velocity magnitudes in WT monolayers were high in the first ∼100-200 µm at the edge and then progressively decreased toward the bulk. In contrast, the velocity magnitudes in KO monolayers were strongly decreased for edge cells and not significantly different from velocities in the bulk **(Figure 2C,D),** indicating that KO cells had lost the ability to migrate faster at the edge than in the bulk of the monolayer. We also extracted from the PIV data a velocity order parameter, here defined as the cosine between velocity vectors and the main direction of the wound edge displacement **(Figure S2E)**. While this parameter, which denotes how the local direction of cell displacements aligned with the monolayer principal direction, was high in WT monolayers, it was strongly decreased in KO monolayers, both at the edge and in the bulk **(Figure 2E,F)**. We also extracted the spatial velocity correlation which defines the distance over which cell velocities remain coordinated **(Figure S2E)**. The correlation of cell velocity vectors was high at short distances up to 100-200 µm for both cell types, however it decreased much faster for KO cells, meaning that these cells are less coordinated at short distances than WT cells. Over 200 µm, a distance of tens of cells, the spatial velocity correlation almost decreased to zero for KO monolayers while it remained significantly higher (∼0.3-0.4) in WT monolayers, **(Figure 2G)**. Altogether, these data show that the coordination of the epithelial monolayer movements was impaired both at short distance and long distance in KO conditions. Since E-cadherin– mediated contacts are essential for coordinated cell movement (Cai et al., 2014; Dumortier et al., 2012; Ladoux & Mège, 2017), we investigated whether E-cadherin expression was altered in these cells. Western blot showed similar E-cadherin levels in both WT and KO cells **(Figure S2B)**, showing that the impaired collective migration observed in vimentin KO cells was not due to a loss of E-cadherin. Altogether, these results show that vimentin expression favors collective migration of epithelial cells by increasing migration speed, order parameter and coordination. To strengthen this hypothesis, we performed a complementary gain of function set of experiments by generating MDCK cells overexpressing vimentin. In similar wound healing assays, in contrast to WT and KO cells, vimentin-GFP overexpressing monolayers displayed strongly increased cell migration speed and spatial velocity correlation **(Figure S2C, D)**. These gain-of-function data support the conclusion that vimentin promotes efficient collective migration.

### Vimentin is required for leader-follower cells coordination and directional progression of fingers

Wound-induced vimentin expression is maximal in fingers **(Figure 1C)** which are polarized mechanical multicellular units essential in guiding the healing monolayer (Reffay et al., 2011, 2014). This led us to investigate in more detail the effect of vimentin loss of function on finger dynamics and mechanics. The absence of vimentin led to delayed formation and aberrant gross shape of fingers, already seen by phase contrast time-lapse imaging **(videos 1,2)**. Tracking of the displacement of the fingertip over 2 hours further showed that the migration path of the finger was more tortuous for KO than for WT monolayers **(Figure 3A)**. We further analyzed the dynamics of cells in well developed (≥100 μm) WT and KO fingers. Vimentin KO-leader cells migrated significantly slower (18 ± 5 µm/h) than WT leader cells (28 ± 11 µm/h) **(Figure 3B)**. Furthermore, KO leader cells as well as follower cells had a significantly lower directional persistence **(Figure 3C, Figure S3A)**, demonstrating that vimentin expression contributes to proper finger dynamics and persistence.

**Figure 3:**
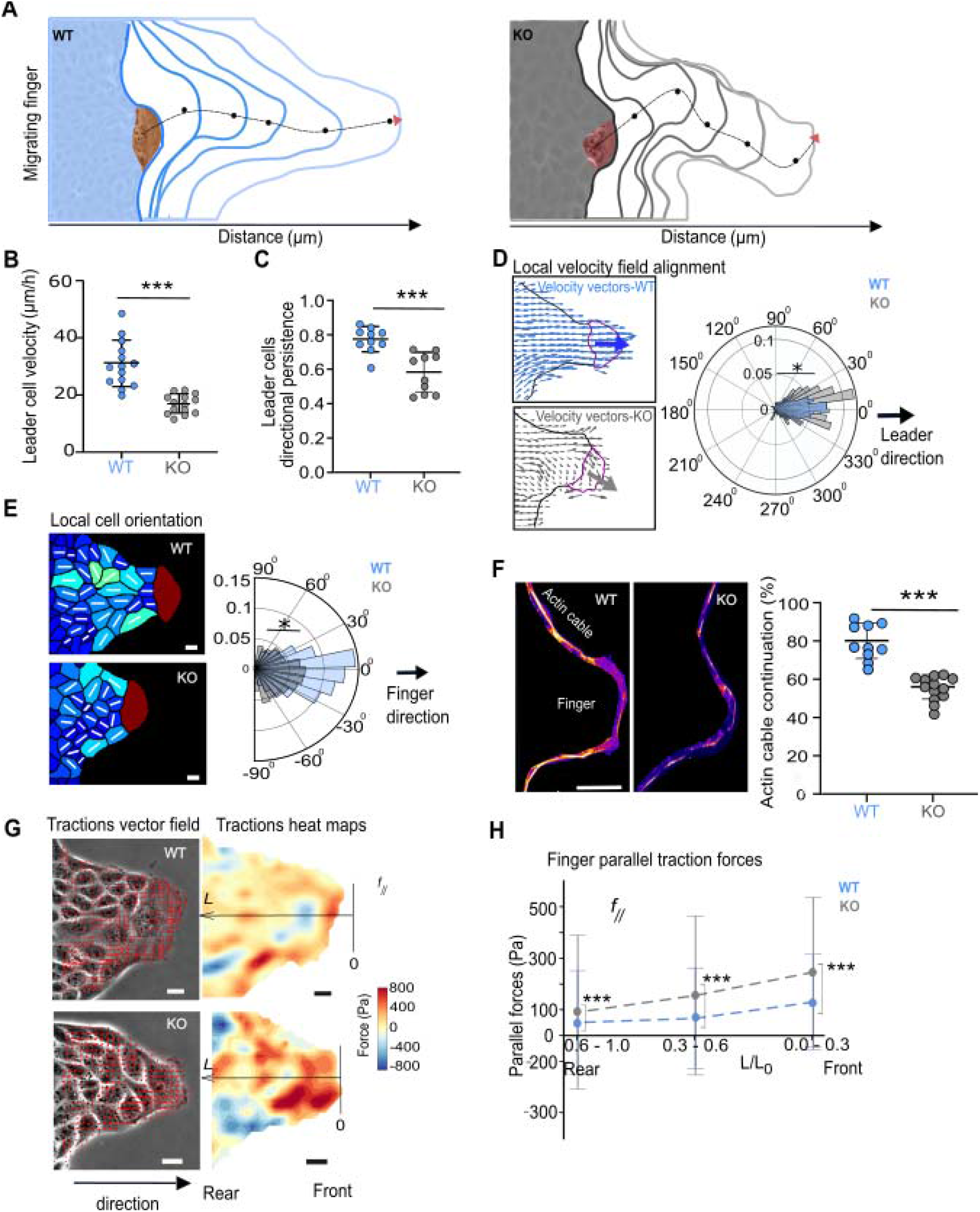
Vimentin loss of function impairs leader-follower cells coordination and directional progression of fingers. **(A)** Tracks of the migrating fingers of time interval 2 hours. Dotted points indicate the centroid of the leader cell over time and black dotted line represents the migration path of the finger. **(B)** Quantification of leader cell velocity, averaged over a two-hour period. Each point represents an individual tracked leader cell. Horizontal lines indicate mean ± SD. Mann-Whitney comparison, n=14 leaders (WT) and n=14 leaders (KO). **(C)** Directional persistence of leaders. Trajectories were analyzed for a 2-hour period, starting from the formation of mature fingers exceeding 100 µm in length. Each point represents an individual tracked leader cell. Horizontal lines indicate mean ± SD. Mann-Whitney comparison, n=10 leaders (WT) and (KO). **(D)** Local velocity order parameter: A schematic depiction of velocity vectors within fingers, indicating leader cells with a magenta boundary. The bold colored arrows represent both the average velocity vectors and the directionality of the leader cell. The polar histogram illustrates the orientation distribution of follower cell velocity vectors relative to the mean velocity vector of the leader cells. Mann-Whitney comparison, n=12 fingers (WT) and n=12 fingers (KO). (**E**) Cell orientation order parameter: Schematic showing cell alignment within the fingers. Vectors (white bars) denote cell major axis orientation. The polar histogram indicates the orientation of the cell relative to the principal axis of the finger (excluded the leader cells in this analysis). **(F)** Immunostaining of actin (Fire LUT) shows the multicellular actin cable that surrounds the finger. The quantification of the actin cable is expressed as a percentage of the total length of the finger. Scale bar, 50 µm. Mann-Whitney comparison, n=9 fingers (WT) and n=13 fingers (KO). **(G)** Mapping of the traction forces in the finger. Phase contrast images of the fingers superimposed with the traction vectors (red), along with the corresponding heat maps of the parallel component (forces opposite to the direction of the finger and parallel to the principal axis) of the traction forces. **(H)** Parallel forces from front to rear of the finger. The data points indicate binned averaged values, colored dotted lines and bars represent the mean ± SD. Mann-Whitney comparisons, n=10 fingers (WT, KO). Sample sizes were representative of three independent experiments.

Fingers are marked by coordinated migration and orientation of their leader cell with follower cells (Reffay et al., 2011). Thus, to better understand the effect of vimentin loss of function on fingers, we analyzed the velocity, shape and ordering of leader and follower cells by a detailed analysis of the PIV-derived velocity fields over the finger area. The averaged velocity vectors of leader and follower cells were extracted from finger PIV maps and their angular orientations compared **(Figure 3D)**. WT follower cells velocity vectors aligned with the average leader cell velocity vector, while velocity vectors of KO follower cells significantly diverged from the main direction imposed by the leader cell. Then, we quantified projected cell area, aspect ratio and major axis orientation. Both leader and follower KO cells showed decreased mean projected cell areas compared to their WT counterparts, with a more pronounced effect for follower cells **(Figure S3B)**. Their aspect ratio was also altered in KO fingers compared to their WT counterparts, with KO cells being significantly less elongated than WT cells **(Figure S3C)**. The main axis of leader cells was oriented perpendicular to the migration axis in both conditions. However, while the major axis of follower cells was preferentially oriented in the direction of the finger, perpendicular to the leader cell major axis, in WT conditions, this ordering was strongly impaired in KO fingers **(Figure 3E)**. These findings reinforce the critical role of vimentin in sustaining the directional progression of fingers, closely linked to cell’s spatial organization and likely to their mechanical coupling. The structural integrity of fingers depends also on the formation of multicellular actomyosin cables connecting follower cells on the side up to the leader cell (Reffay et al., 2014). F-actin staining revealed that the continuity of the multicellular actin cable along the edge of the finger was significantly disrupted in KO conditions compared to WT **(Figure 3F)**. In WT fingers, the actin cable spanned 80 ± 10% of the edge of the finger, while in KO it was reduced to 55 ± 5%. Vimentin expression is thus also necessary to maintain contractile multicellular actin cables alongside the fingers, enabling these protrusions to function as a coherent, mechanically integrated structure.

Therefore, we performed Traction Force Microscopy (TFM) on the whole monolayers **(Figure S4A, B)**. Interestingly, the traction forces magnitudes were overall increased in the KO monolayers, consistently with previous reports (Costigliola et al., 2017; De Pascalis et al., 2018). We further analyzed the TFM data in mature fingers **(Figure 3G, S4C)**, then extracted the spatial distribution across the fingers of the forces perpendicular (f_┴_) and parallel (f_‖_) to the direction of finger progression. Perpendicular forces increased from the main axis to the outer edges of the finger, with a marked increase in magnitude in KO fingers **(Figure S4C, D)**. Parallel forces decreased from the front to the back of the finger in both conditions but were significantly higher along KO finger length compared to WT **(Figure 3H)**, indicating that vimentin expression in fingers contributed to restrict perpendicular as well as parallel forces along the finger length. Together, these results demonstrated that the wound-induced expression of vimentin contributes to the supracellular organization and coordination of leader-follower cells, the directional progression of fingers, and simultaneously reduces traction forces exerted by the cells on the substratum.

### Vimentin contributes to the organization of traction force and focal adhesion distributions in leader cells

Leader cells, which are found at the tip of multicellular fingers, are the cells exerting the highest traction forces on the substratum **(Figure 3G, H)**. We thus further studied how the loss of function of vimentin impacts their mechanics. Traction force maps showed an overall increase in traction forces under KO compared to WT leader cells **(Figure 4A)**. The magnitude of traction force applied by KO cells was almost double to the one of WT cells **(Figure 4B)**. Both the Tx (parallel) and Ty (perpendicular) components of these traction forces followed the same trend **(Figure S4E, G)**. Quantitative analysis of the spatial distribution of traction forces from the front to the rear of leader cells revealed a non-monotonous profile in WT cells with a peak of force a little away from the front edge of the cell, then a decrease in force magnitude toward the back of the leader cell **(Figure 4C)**. This was similarly observed for Tx, Ty force components **(Figure S4F, H)**. In contrast, in KO leader cells, we observed a sharp increase in force magnitudes, as well as Tx, Ty, from the cell front to the back, rapidly reaching a plateau, without further decrease when going toward the cell rear **(Figure 4C, S4F, H)**. Thus, KO leader cells developed much higher forces than WT cells, these forces were more evenly distributed and in particular high under their cell body. This suggests that vimentin expression is critical for modulating both the magnitude and the subcellular distribution of traction forces in leader cells.

**Figure 4:**
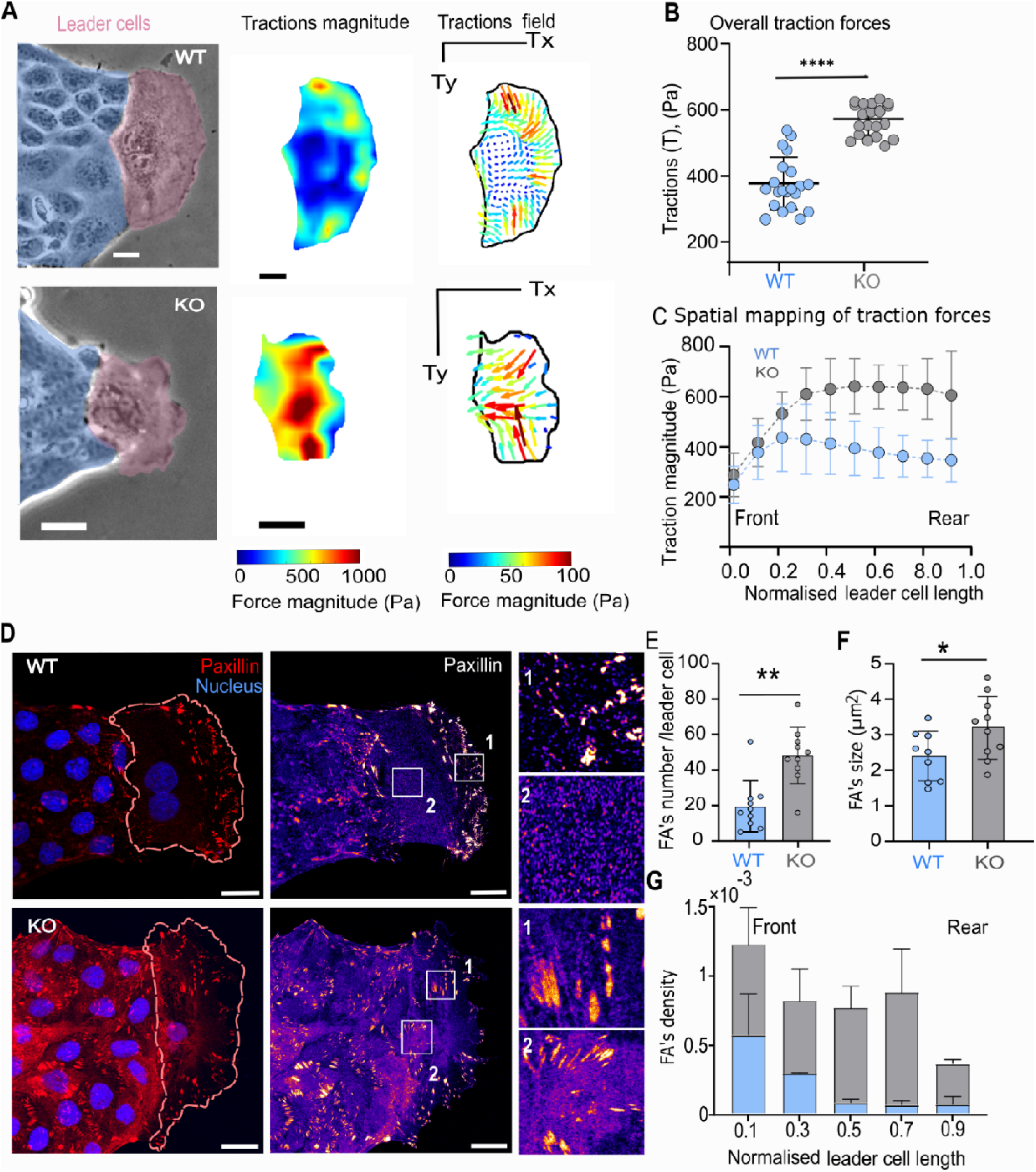
Vimentin loss of function induces the redistribution of traction forces and focal adhesions in leader cells. **(A)** Phase contrast images of migrating leader cells (pink). Scale bars: 20 µm. The corresponding heat maps show the magnitude and traction vectors. **(B)** Quantification of tractions for every 10 min for 2 hours calculated by averaging the magnitude of forces detected under each leader cell at each time point throughout the observation period. Horizontal lines indicate mean ± SD. Mann-Whitney comparisons, n=21 leaders (WT) and n=19 leaders (KO). **(C)** Quantification of spatial distribution of tractions in leader cells from front (right) to rear (left). The data points represent mean ± SD, binned into 0.1-unit intervals. **(D)** Images of migrating leader cells stained for paxillin (red), and nucleus (blue). The fire LUT images illustrate the focal adhesions distribution. Enlarged images of focal adhesions at the front and rear of the leader cell are shown in the inserts indicated with front (1) and rear (2). **(E, F)** Quantification of focal adhesions (FAs) number and size. Each data point represents the focal adhesion number and area per leader cell. Bars indicate mean ± SD. Mann-Whitney comparisons, FA’s number and size n=9 leaders (WT), n=10 leaders (KO). **(G)** Quantification of the focal adhesion density along the normalized length of the leader cell from front (right) to rear (left). FA density was calculated as the number of adhesions per unit cell area in each binned segment. Bars indicate mean ± SD. Sample sizes were representative of three independent experiments.

The alterations in both the magnitude and distribution of traction forces observed in leader cells in the absence of vimentin expression may reflect a redistribution of FAs. Both the number and size of FAs as revealed by paxillin immunostaining were significantly increased in KO compared to WT leader cells (**Figure 4D-F)**. In addition, whereas FAs were more present at the front of WT leader cells, they were almost evenly distributed from the front to the rear of KO leader cells **(Figure 4G)**. These findings unraveled that vimentin expression strongly regulates the quantity, size, and spatial distribution of FAs in leader cells. The perturbed distribution of traction forces and FAs in KO cells suggested that the actomyosin networks could thus be disorganized in the absence of vimentin. Accordingly, in KO monolayers, stress fibers and their associated focal adhesions were markedly enhanced in fingers (**Figure S4I**), suggesting that the altered distributions of stress fibers and FAs in KO cells is maintained deep in the monolayer. Altogether, these results show that vimentin expression in migrating leader cells and fingers is critical in maintaining the spatial restriction of FAs, stress fiber and associated traction forces at the front of the monolayer.

### Vimentin contributes to the organization and dynamics of actin arcs in leader cells

The actin cytoskeleton organization in branched actin networks and actin arcs plays a critical role in driving cell protrusive activity (Burnette et al., 2011; Koestler et al., 2008; Svitkina, 2007). As the coordination of the two actin modules is required for both the formation of FAs and cell migration (Burnette et al., 2011; Koestler et al., 2008; Kumari et al., 2024; Yamaguchi & Knaut, 2022) we investigated their distribution in KO leader cells **(Figure 5A)**. Double vimentin/F-actin immunostaining revealed that vimentin filaments were restrained at the back of actin arcs of WT leader cells **(Figure 5A, B)**, as reported in single U2OS cells (Jiu et al., 2015). KO leader cells showed disrupted and bundled actin cables in the lamella instead of well-defined actin arcs observed in WT cells **(Figure 5A)**. The overall F-actin staining, predominant in the lamella in WT cells was spread within the cell body in KO cells **(Figure 5B)**, The transverse F-actin bundles were positioned more centrally in KO cells, distant ∼12 µm from the lamellipodia tip, compared to ∼7 µm in WT cells **(Figure 5C)**, and their thickness was also increased (**Figure S5A**). Moreover, live cell imaging revealed that their dynamics was also affected in KO leader cells. Indeed, kymograph revealed significantly slower retrograde movement of these actin cables in KO leader cells compared to WT cells **(Figure 5D, S5B, video5)**.

**Figure 5:**
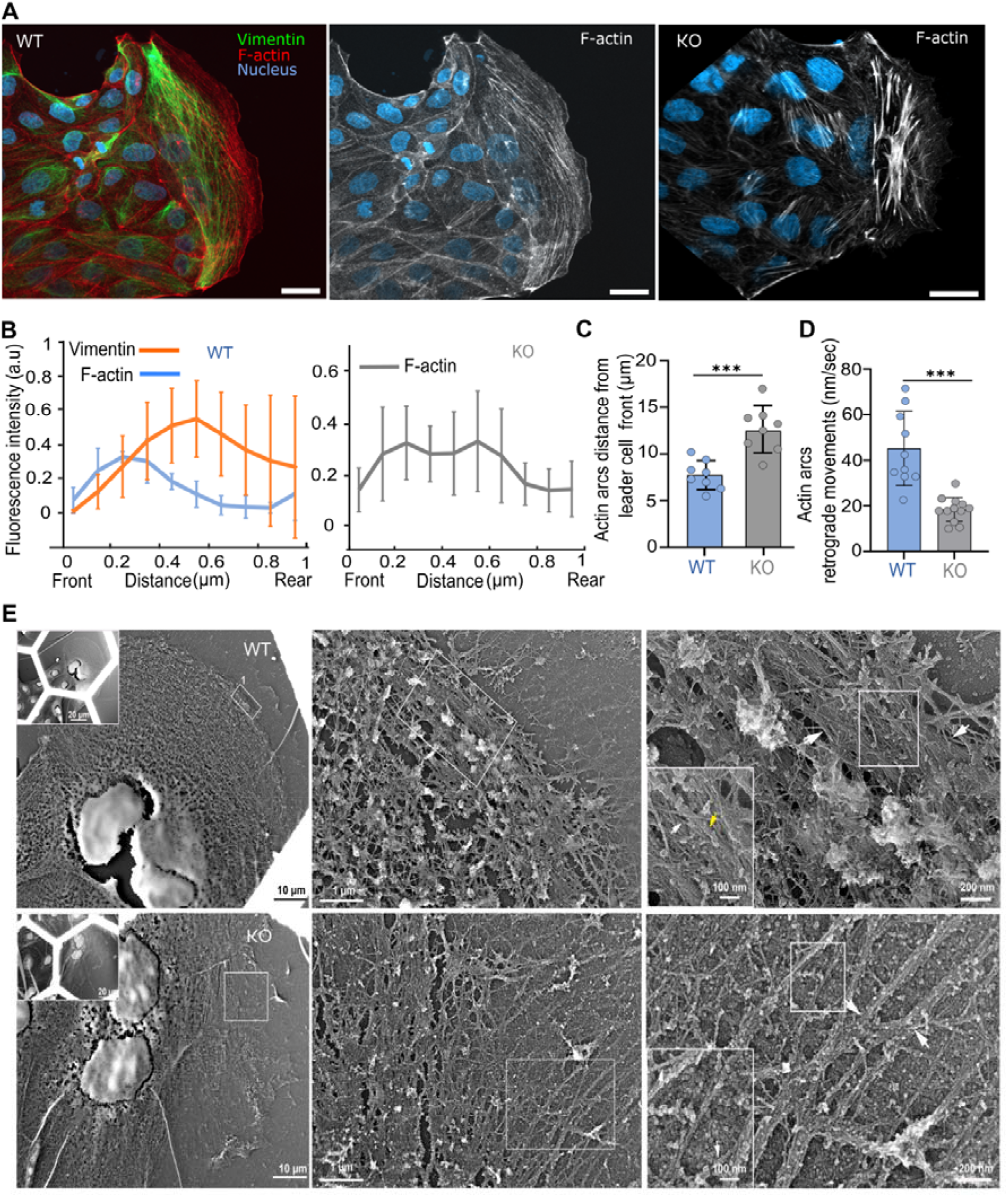
Vimentin loss of function disorganizes the contractile actin arc in leader cells. **(A)** Immunofluorescence images showing the co-distribution of vimentin (green), F-actin (red), and nuclei (blue) in migrating leader cells. Gray scale panels: F-actin staining in WT and KO leader cells. Scale bars, 50 µm. **(B)** Quantification of fluorescence intensity profiles of vimentin (orange) and F-actin (blue) in WT and F-actin (gray) in KO leader cells. The colored thick lines and colored bars indicate mean values ± SDs. Mann-Whitney comparisons for cell area n=8 leaders (WT) and n=6 leaders (KO). **(C)** Quantification of actin arcs distance from the lamellipodia front edge. Bars indicate mean ± SD. Mann-Whitney comparisons, n=8 leaders (WT), n=8 leaders (KO). **(D)** Quantification of actin arcs retrograde movements. Bars indicate mean ± SD. Mann-Whitney comparisons, n=8 leaders (WT), n=8 leaders (KO). **(E)** Electron microscopy images of actin arcs in WT and KO leader cells at progressive magnifications. Left: Low magnification showing overall structure; Right: higher magnification insets highlighting the actin arcs organization. White arrows indicate actin arcs, and yellow arrows mark microtubules. Sample sizes were representative of three independent experiments.

To further investigate the actin organization at the leading front, we used high-resolution electron microscopy using platinum-replica electron microscopy (PREM) **(Figure 5E)**. WT cells displayed well-defined and spatially ordered actin arcs aligned precisely at the lamellipodia-lamella interface, forming a coherent supracellular structure consistent with directed lamellipodia protrusions. In contrast, KO cells exhibited severely disrupted arc organization, with actin bundles forming irregular, fragmented structures that were dislocated away from the leading edge. This loss of spatial confinement of actin arcs indicates a fundamental defect in cytoskeleton self-organization, that may directly impair protrusive stability and front-rear polarization. We hypothesized that vimentin’s role in actin remodeling within protrusions could be observed at the single-cell level. Consistently, similar actin organization was seen during isolated cell spreading **(Figure S5C)**. 3-4 hours after seeding, two populations of WT cells emerged, one had a non-polarized, isotropic shape while the other had already acquired a polarized shape. Non-polarized WT cells showed a continuous circumferential actin arc in the lamella, while polarized WT cells had undergone actin symmetry breaking and displayed a well-organized actin arc at the lamellipodia rear **(Figure S5C)**. KO cells also displayed both polarized and non-polarized populations. However, non-polarized cells exhibited a disrupted perinuclear F-actin ring, shifted closer to the nucleus, from which numerous thicker and longer radial actin cables emerged **(Figure S5C)**. Some KO cells could still polarize but the actin arcs at the back of the lamellipodium were disorganized and shifted toward the nucleus, mirroring the observations made in KO leader cells. These data highlight that mechanical cooperation between vimentin and actin cytoskeletons is essential for maintaining actin arcs restricted to the lamella-lamellipodia interface.

### Vimentin controls the dynamics and persistence of the lamellipodium of leader cells

Actin arcs in the lamella serve as structural elements underlying the connection between the lamella and the lamellipodium during epithelial cell directed motion (Burnette et al., 2011). Their disorganization in KO leader cells may thus impact lamellipodia dynamics. We thus analyzed the lamellipodia dynamics in leader cells by time-lapse DIC imaging **(Figure 6A, video 6)**. Kymograph analysis revealed an overall decrease in the velocity of both the protrusions and the retractions of the lamellipodia of KO leader cells compared to WT **(Figure 6B)**. Moreover, we observed a significant decrease in protrusion frequencies in KO cells **(Figure 6C)**. Interestingly, retractions lasted significantly longer than protrusions in KO lamellipodia (**Figure 6D**). Altogether, these data reveal an altered dynamics of lamellipodia that could lead to an overall lower ability of the KO leader cell to perform efficient protrusion. Accordingly, we observed a delay in the appearance of leader cells in wounded KO monolayers **(Figure 6E)**. Moreover, the appearance of lamellipodia in a few edge cells, which preceded leader cell appearance, was also delayed and significantly decreased in KO monolayers compared to WT monolayers (**Figure 6F)**. This was confirmed by the delay in spreading and polarization observed in isolated KO cells **(Figure S6A-C)**. These observations suggest that vimentin expression at the wound edge favors lamellipodia appearance in a cell autonomous manner.

**Figure 6:**
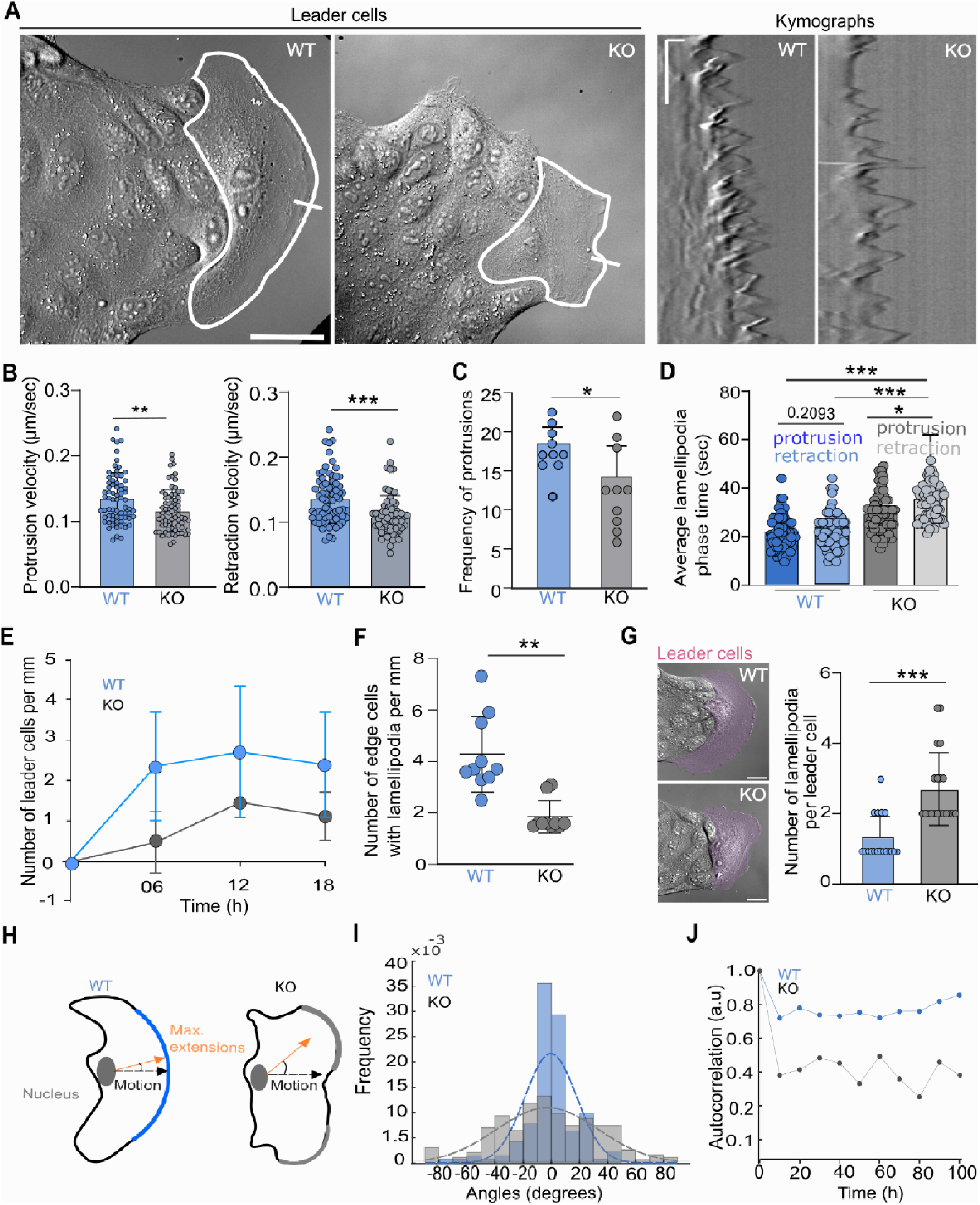
Vimentin loss of function impairs the dynamics of lamellipodia in leader cells. **(A)** Representative Differential phase contrast (DIC) Images of migrating leader cells, with segmented cell outlines. Kymographs (right) illustrate edge dynamics over time for each condition. The regions of interest marked by white lines for kymograph analysis. Images were acquired every 2 seconds for 10 minutes. Scale bars, 50 μm (DIC images), 2 μm (horizontal) x 120 sec (vertical) in kymographs. **(B)** Quantification of protrusion and retraction velocities. Bars indicate mean ± SD. Mann-Whitney comparison, n=10 leaders, (WT) and n=10 leaders (KO). **(C)** Frequency of protrusions and retractions, mean ± SDs. Mann-Whitney comparisons, n=10 leaders (WT) and n=10 leaders (KO). **(D)** Average duration of lamellipodia phase events per unit time. Bars indicate mean ± SD. Mann-Whitney comparisons for cell area n=10 leaders (WT) and n=10 leaders (KO)**. (E)** Quantification of leader cell density over time in the migrating monolayers. The density of leader cells was normalized relative to the width of the monolayer leading edge. Data points and bars indicate mean ± SD. Mann-Whitney comparison, n=10 leaders (WT), n=10 leaders (KO). **(F)** Quantification of the number of edge cells with a lamellipodia at time 6h. Each data point indicates leader cell. Bars indicate mean ± SD. Mann-Whitney comparison, n=10 leaders (WT), n=10 leaders (KO). **(G)** Representative DIC images indicate leader cells (pink). Scale bars, 20 µm. The corresponding graph represents average number of lamellipodia formed in the leader cell over time. Bars indicate mean ± SD. Mann-Whitney comparison, n=10 leaders, (WT) and n=10 leaders (KO). **(H)** Diagram illustrating the angular orientations of lamellipodial extension relative to the direction of cell migration. The orange and black arrows denote the directions of lamellipodia new extensions (thick borders in blue and gray) and cell movement, respectively. **(I)** The corresponding histogram illustrates the distribution of lamellipodia protrusion orientations. Sample sizes n=10 (WT) and n=10 (KO) **(J**) Directional autocorrelation analysis showed that lamellipodia extensions are less persistent, as evidenced by the faster decay of their autocorrelation curve (since slower decay reflects higher directional persistence). Sample sizes n=10 (WT) and n=10 (KO). Sample sizes were representative of three independent experiments.

Then, we addressed how vimentin loss of function impacted lamellipodial persistence on a larger time scale in leader cells by following by phase contrast imaging the evolution over time of leader cell lamellipodia shape **(video 7)**. Indeed, lamellipodia persistence has been linked to lamellipodia dynamics (Yolland et al., 2019). Thus, we quantified the bulging of new lamellipodial extension in leader cells and found a significant increase in KO leader cells compared to their WT counterparts **(Figure 6G)**. Moreover, the angular orientations of these protrusions which peaked in the direction of the leader cell migration for WT cells, significantly spread at higher degrees for KO cells **(Figure 6H, I)**. Directional autocorrelation analysis revealed that the directional persistence of the lamellipodia over time was significantly decreased in KO cells **(Figure 6J)**. To further explore how vimentin loss of function might affect collective cell behavior through disrupted front-rear polarization, we employed a previously described model system of cells migrating in constrained rings where cells collectively polarize at the level of single cell to support coordinated migration (Jain et al., 2020). In this system, upon reaching confluency, WT cells acquire coordinated front-rear polarization as reported previously, while KO cells fail to coordinate, showing oscillatory movements instead of persistent rotational behavior **(Figure S6D, E, video 9)**. This suggests that vimentin plays a key role in acquiring and maintaining a sustained directional front-to-rear polarity.

To further support this hypothesis we addressed whether cryptic lamellipodia associated to coordinated front-rear polarization described in (Jain et al., 2020) and others 2D (Ozawa et al., 2020) and 3D contexts (Williams et al., 2022) were affected by the loss of function of vimentin. To do so we performed wound healing experiments using mixed populations of MDCK cells (WT or KO) containing a small fraction of the same cells expressing YFP-PBD, a reporter of Rac activity, as a proxy of lamellipodia and cryptic lamellipodia (Jain et al., 2020). We then analysed by live imaging the distribution of Rac activity in leader and follower cells as well as cells more in the back, analyzing the evolution of the shape of the cells, of the protrusions/retraction along their contour as described in (Marshall-Burghardt et al., 2024) **(Figure 7)**, as well as of the accumulation of the YFP-PBD signal along the cell contour **(Figure S7)**. While the WT cells whatever their position showed smooth and persistent lamellipodia and cryptic lamellipodia **(Figure 7A, Videos 10, 11)** associated with persistent polarized distribution of protrusions and retractions **(Figure 7B)**, lamellipodia were more convoluted for KO cells **(Figure 7C, Videos 12-14))**, associated with a significant decrease in protrusion width and persistence **(Figure 7D-F)**. These observations thus confirm that the loss of function of vimentin alters lamellipodia persistence and polarity both in edge cells and in cells behind in accordance with the lack of coordinated front-rear polarization.

**Figure 7:**
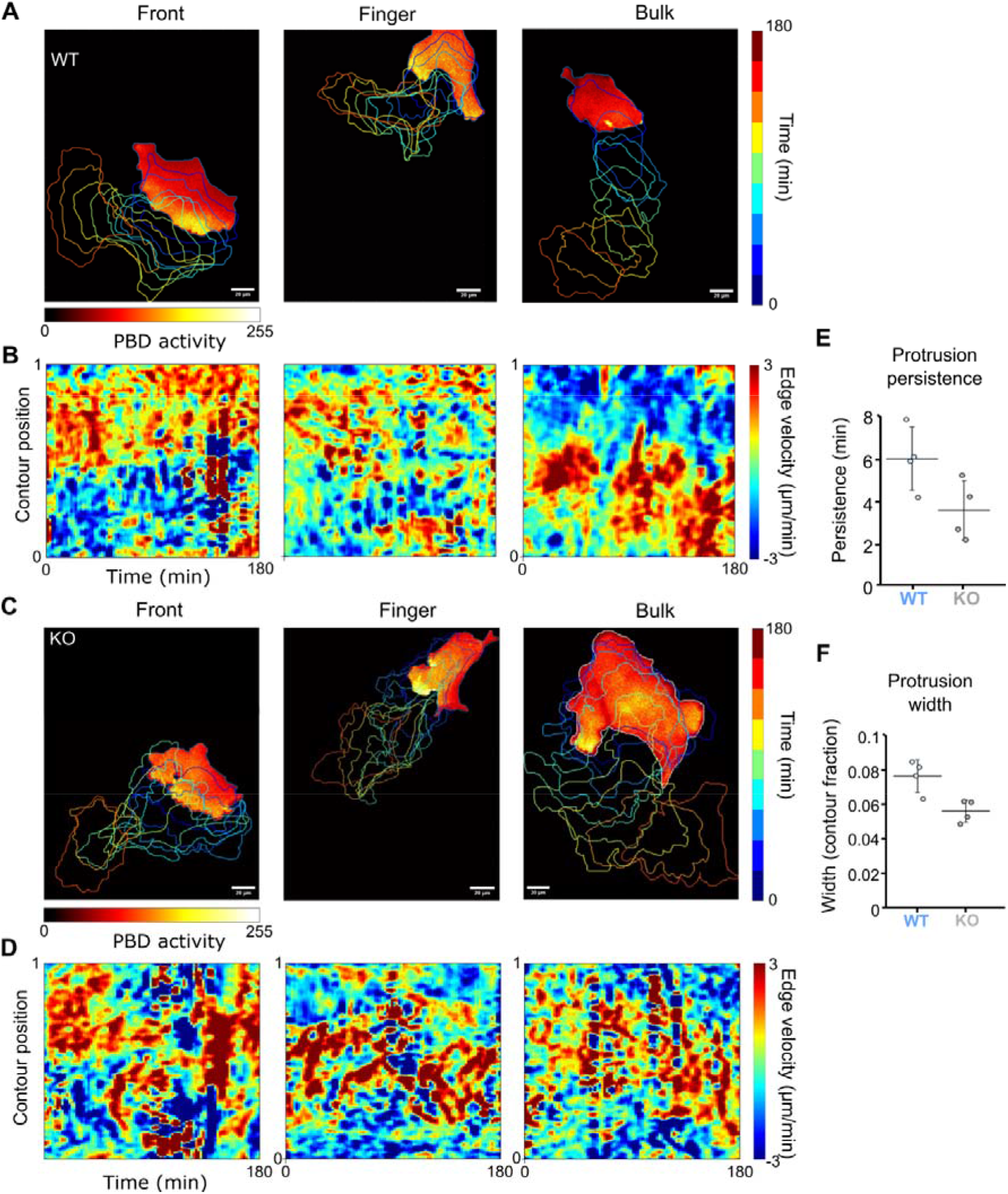
Vimentin loss of function impairs the dynamics and persistence of lamellipodia in both front, finger and bulk cells. **(A)** PBD-YFP signal in front, finger and bulk cells of a WT migrating epithelial monolayer and outlines over time. **(B)** Edge velocity maps along time of the corresponding cells. Negative velocity indicates retraction while positive marks protrusions. X-axis represents time (interval 2 min) and Y-axis represents the linearised cell contour, from 0 (beginning) to 1 (end after complete rotation around the cell’s contour), arbitrary units. **(C)** PBD-YFP signal in front, finger and bulk vimKO cells. **(D)** Edge velocity maps of corresponding Vim KO cells. **(E)** Mean protrusion persistence in WT and Vim KO cells. **(F)** Mean protrusion width (contour fraction) in WT and Vim KO cells. Representative examples out of 3 cells analysed for each condition.

## Discussion

Our findings show that vimentin expression is highly upregulated at the edge of healing epithelial monolayers, in decreasing gradient from the leading edge towards the bulk of the monolayer, with the highest levels in leader cells. Our observations suggest that the exposure of cell assemblies to a free-edge either in wounded monolayers, in sparse cultures or for cells on rings is sufficient to induce vimentin, although we do not know by which signaling mechanisms. However later, vimentin accumulates preferentially in fingers rather than remaining uniformly distributed along the wound boundary. We further show that vimentin expression positively regulates collective migration by promoting both the cell autonomous initiation of motility and the coordination of cell ensembles. Notice that the stimulating effect of vimentin expression on the cell motility and coordination is supported by the observed opposite effects of loss and gain of function. Vimentin is required for enhanced cell coordination and force balance both in the bulk and at the edge of the monolayer. We further demonstrate that elevated vimentin levels in finger structures supports directed migration of the entire cell cohort. Vimentin regulates leader-follower cell coordination and mechanical coupling, which are critical for maintaining directional migration **(Figure 8)**. In leader cells, vimentin is required for the polarization of focal adhesion and traction force distribution, as well as for lamellipodia dynamics required for efficient and directed migration. Vimentin does so by stabilizing actin arcs at the lamella-lamellipodium interface, which is essential for coupling force generation with mechanical polarization (Burnette et al., 2011). The fact that the defects observed on collective cell migration are associated with strong alteration of the actin cytoskeleton and lamellipodia dynamics in the absence of cell-cell contacts, in single cells, indicates that the defects of collective cell induced by vimentin loss of function are primarily cell autonomous. Although, E-cadherin levels remain unchanged in vimentin KO cells, and no major distribution of desmoplakin (data not shown) were detected we cannot fully exclude an indirect effect through subtle alterations of adherens junctions or desmosomes.

**Figure 8.**
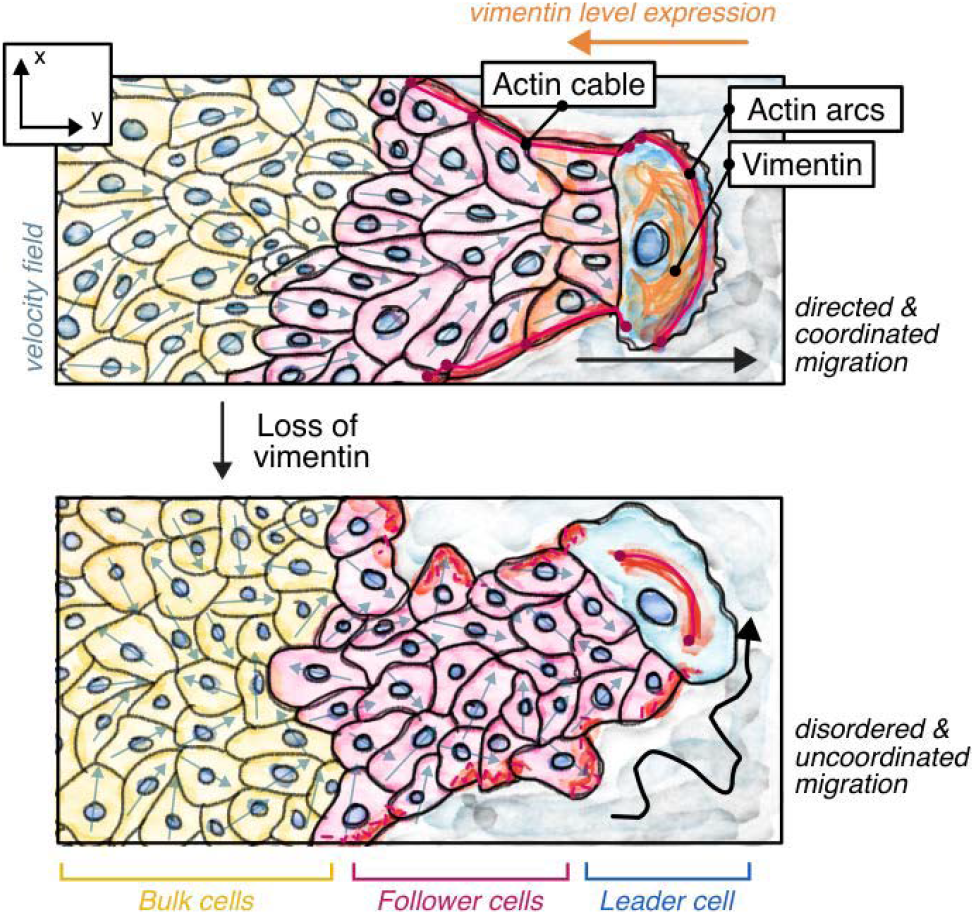
Proposed model for the role of vimentin in epithelial collective migration. In WT conditions, vimentin expression at the edge of the monolayer supports correct organization and polarization of the leader cell, as well as the stability and directionality of its lamellipodia, leader-follower alignment, coherent finger organization and long range coordination of cell migration across the monolayer (Top). In vimentin KO monolayers, leader cell internal organization and dynamics, in particular of the lamellipodia and actin arcs, as well as leader – follower cells coordination are impaired, resulting in slower and less coordinated migration (Bottom).

Our studies bring together observations reported previously in other cell types: in the absence of vimentin, monolayers collectively move less efficiently. This is related to overall less persistent, less coordinated motion, and ultimately less directed motion at the scales of both the monolayer and epithelial fingers (De Pascalis et al., 2018). On a mechanical side, this correlates to a disorganization of the spatial distribution of traction forces and focal adhesions, which are no longer polarized and restricted to the tip of fingers and leader cells, an overall increase of traction force magnitude and a reinforcement of stress fibers and focal adhesions (Costigliola et al., 2017; De Pascalis et al., 2018; Jiu et al., 2017). The latter features likely stem from the destabilization of lamellar actin arcs previously reported to be supported by the vimentin-actin arc interplay (Jiu et al., 2017). This brings out the question of the connection between these three major facts, which one can reformulate as: (i) how does this morphological change of actin arcs affects cell polarity and migration? (ii) how do those cell-scale events dysregulate collective cell migration? Our data put answers forward to both questions.

First, zooming in to the cell and lamellipodium scales, we observed the effects of vimentin depletion on cell migration dynamics at various time scales: on the minute scale, it makes the protrusion-retraction cycles of the lamellipodia longer-lived; on the hour scale, the same lamellipodia are more split along the cell edge, and their angular orientation is widened; on the scale of tens of hours, cells tend to remain less polarized and migratory, be it as isolated cells or as cell groups in closed rings. Altogether, those facts suggest that by restricting the actin arcs close to the lamellipodium, vimentin ensures a good connection between the backward flowing branched actin and the contractile arcs that collects it (Burnette et al., 2011). In the absence of vimentin, this connection is loosened so that lamellipodia would become floppy, hence reducing their directional persistence and the overall cell polarization. It is remarkable to notice that these processes are fully cell autonomous and affect in the absence of vimentin both the lamellipodia of edge cells as cells and cryptic lamellipodia of cells behind.

Then, zooming back out to the collective scale, how could those local changes translate into defective collective migration? Our wound healing and closed rings data could be explained by two – non-exclusive – hypotheses: (i) an explicit alignment interaction mechanism, that directly couples the velocity alignment of a cell to that of its neighbor, as proposed in a theoretical work by Nir Gov team (Tarle et al., 2015), which is missing in the absence of vimentin. Notice that this theoretical work predicts an essential role of both velocity alignment interactions and intercellular contractile actomyosin cables in finger formation and stability, in the case of wound healing; two parameters strongly affected by vimentin loss of function in the present work. (ii) A strong cell polarity alignment affected by vimentin loss of function. This may result from an impact of unbalanced cell-substratum and cell-cell adhesion forces on self-reinforcing cell polarity mechanisms. This process may involve the contribution of the propagation of ERK waves that have been reported to regulate collective cell migration and collective cell polarization in many epithelial cell models (Aoki et al., 2017; Hino et al., 2020; Matsubayashi et al., 2004). Although we do not exclude the two hypotheses, especially in the fingers where the multicellular lateral actin cable is disrupted, our data point to hypothesis (ii). Indeed, upon vimentin depletion, a cell fails to self-polarize when it is isolated or flanked by two equally non-moving cells; on the contrary, it orients reasonably well on average, although not efficiently, when the monolayer is subjected to a global directional cue.

Recent works report that loss of vimentin significantly impairs the ability of spheroids to collectively expand in collagen gels (Thanh et al., 2024; Van Bodegraven et al., 2025) suggesting that vimentin is also required to promote collective cell migration in 3D. Thus, our findings are also relevant for 3D collective migration although this migration is more complex and associated with MMP-dependent degradation and remodeling of the matrix which is not possible in our 2D system.

In conclusion, our study reveals that vimentin regulates the emergent mechanical forces of epithelial monolayers undergoing partial EMT. By regulating active contractile forces, and intercellular mechanical coupling, vimentin controls the speed, directionality, and coherence of collective cell migration across multiple scales from individual cell motility to coordinated finger like protrusions and large-scale monolayer movements. Loss of vimentin function disrupts mechanical coordination and force distribution at the leading edge, impairing protrusive stability and collective guidance. These findings identify vimentin as a central mechanical integrator linking actomyosin generated forces to collective tissue dynamics. Further studies are needed to evaluate whether this key marker of epithelial-mesenchymal transition also contributes by this mechanism to cancer cell migration out of carcinoma and initiation of invasion in surrounding tissues.

## Materials and Methods

### Cell culture

MDCK WT cells (ATCC CCL-34) and MDCK vimentin KO cells were cultured in DMEM medium (GlutaMAX, high glucose, and pyruvate; Life Technologies) supplemented with 10% fetal bovine serum (FBS; Life Technologies) and 1% penicillin-streptomycin (Life Technologies). Cultures were kept at 37°C in a humidified incubator with 5% CO. To maintain a consistent fluency of 70%, cells were passed 2-3 times per week.

### Generation of vimentin knockout and vimentin-GFP overexpressing MDCK cell lines

MDCK vimentin knockout stable cells (KO) were obtained using a CRISPR-Cas9 double nickase plasmid from Santa Cruz Biotechnology, containing a pool of 3 different gRNA plasmids: TGTCGCGCTCCACCTCGACG, CGGCTGTTGCAGGTTGCAAG and TCCTCCTACCGCAGGATGTT. Approximately 2 million cells were electroporated using an Invitrogen Neon Transfection System to develop stable transfectants. The electroporation conditions were a single 20 ms pulse at 1650 V with 3 µg of plasmid DNA. After electroporation, cells were incubated for 24 hours before adding 2.5 µg/ml puromycin to the medium for selection, allowing only successfully transfected cells to survive. Forty-eight hours after electroporation, single GFP-positive cells were sorted into 96-well plates using flow cytometry with a BD Biosciences Influx 500 sorter analyzer. Clones were selected based on the absence of vimentin protein by immunofluorescent staining. The absence of vimentin in the clones generated was confirmed by analysis of protein extracts using western blot analysis (Figure S2A). Vimentin-GFP overexpressing cells were generated by electroporating a mouse vimentin GFP cDNA in a pEGFP vector (Mignot et al., 2007) and then selected under geneticin. WT MDCK cells expressing YFP-PBD were described previously (Jain et al., 2020). The YFP-PBD plasmid (obtained from Dr. Fernando Martin Belmonte, Universidad Autónoma de Madrid, Spain) was electroporated in Vim KO cells, then selected under geneticin to obtain YFP-PBD Vim KO cells.

### Electrophoresis and Western blotting

Cells were grown in tissue culture dishes until full or partial confluence. Cells were lysed at 4°C for 20 minutes using a buffer containing 10 mM Tris (pH 7.5), 150 mM NaCl, 0.5% NP 40, 0.5% Triton-X 100, 10% glycerol, 1x protease inhibitor cocktail (Roche), and 1x phosphatase inhibitor (Phosphostop, Roche). Lysates were centrifuged at 13,000 g for 15 minutes, and protein concentration was determined using the Bradford assay (BioRad). Proteins were separated by SDS-PAGE on a 4-12% Bis-Tris gel, transferred to a nitrocellulose membrane, and blocked with 5% nonfat dry milk in PBS with 0.1% Tween 20. Membranes were incubated overnight at 4°C with primary antibodies (1:1000), followed by secondary HRP-conjugated antibodies (1:1000). Immune complexes were detected using SuperSignal West Femto substrate and visualized with a ChemiDoc system (BioRad). Band intensities were quantified using ImageJ, with GAPDH as a loading control for normalization.

### Immunofluorescent Staining

The confluent and migrating monolayers were fixed and permeabilized simultaneously with a pre-warmed PFA solution (2.5% PFA and 0.1% Triton X-100) for 10 minutes. To block nonspecific binding, the samples were treated with 1% bovine serum albumin (BSA) in phosphate-buffered saline (PBS) for one hour. Primary antibodies were then added, and the samples were incubated overnight in the same 1% BSA solution. Subsequently, the samples were incubated with secondary antibodies for 2 hours. The nuclei were stained with DAPI or Hoechst blue and mounted with Mowiol. Confocal images were captured using a Zeiss LSM 780 and 980 inverted microscopes. The primary and secondary antibodies used, along with their respective concentrations and the manufacturer are listed in **Table 1**.

**Table 1:** Primary antibodies used in this study.

| **Antibodies** | **Species** | **Source** | **Reference** | **Dilution** |
| --- | --- | --- | --- | --- |
| Vimentin | Mouse | Abcam | ab8978 | 1:100 |
| Phalloidin 568 | - | Life Technologies | A34055 | 1:200 |
| Phalloidin 647 | - | Life Technologies | A22287 | 1:200 |
| Alpha-tubulin | Rat | Bio-Rad | MCA77G | 1:100 |
| Cytokeratins | Guinea Pig | Biovalley | GP11 | 1:100 |
| E-cadherins | Rat | Takara | ECCD2 | 1:100 |
| Paxillin | Rabbit | Abcam | ab32084 | 1:100 |
| DRAQ5 | - | Abcam | ab108410 | 1:1000 |

### Electron microscopy

Adherent plasma membranes from cultured MDCK WT cells and MDCK vimentin KO cells grown on glass coverslips were detergent-extracted. Cells were treated with extraction buffer (2 mL stock buffer (5X Stock Buffer: 500 mM 1,4-piperazinediethanesulfonic acid, 25 mM ethylene glycol tetra acetic acid, 25 mM MgCl2, pH’d and kept at 4C), 4 mL 10% PEG (35,000 MW), 4 mL milliQ H2O, 100 uL of TritonX-100, 10 uM nocodazole, and 10 uM phalloidin) for 30 min, followed by a 1 min wash with wash buffer (2 mL stock buffer, 8 mL milliQ H2O, 10 uM nocodazole, 10 uM phalloidin), followed by fixation (2% PFA, 2% glutaraldehyde) for 20 min. Extracted cells were further sequentially treated with 0.5% OsO4, 1% tannic acid, and 1% uranyl acetate before graded ethanol dehydration and hexamethyldisilazane (HMDS) substitution (LFG Distribution, France). Dried samples were then rotary shadowed with 2 nm of platinum (sputtering) and 4-6 nm of carbon (carbon thread evaporation) using an ACE600 metal coater (Leica Microsystems, Germany). The resulting platinum replica was removed from the glass using 5% hydrofluoric acid, thoroughly washed in distilled water, and mounted on 200-mesh formvar/carbon-coated EM grids. The grids were mounted in a eucentric side-entry goniometer stage of a transmission electron microscope operated at 120 kV (JEOL, Japan). The images were adjusted for brightness and contrast using Adobe Photoshop (Adobe, USA) and presented in inverted contrast.

### Migration experiments

For experiments involving monolayer migration, Petri dishes containing a culture insert (culture insert 2-well in µ-Dish 35 mm, Ibidi, Germany) were used. Alternatively, custom-made PDMS blocks, pretreated with a 0.2% pluronics solution for 45 minutes, were employed in culture dishes coated with fibronectin. Cells were seeded in 2 ml of DMEM around the culture insert or PDMS block. Following an overnight incubation period to ensure the formation of a confluent monolayer, the insert was carefully removed, and the area was gently rinsed with DMEM to eliminate nonadherent cells. Subsequently, 2-3 ml of fresh DMEM medium was added to the culture plate before transfer to Biostation IM-Q (Nikon) at 37°C with 5% CO□. The samples were then incubated in a Biostation IM-Q (Nikon) at 37°C with 5% CO□. Live phase contrast imaging was captured every 10 minutes for a total of 24 hours using a 10X objective.

### Live cell imaging

For migration experiments, phase-contrast images were captured every 10 minutes on the Biostation IM-Q (Nikon) at 37°C with 5% CO□. For traction force microscopy (TFM) experiments, both phase-contrast images and fluorescent bead signals were acquired under the same temperature and CO□ conditions. Live imaging of cytoskeletal and Rac sensor dynamics was performed using 60X or 40X objectives on a spinning disk confocal microscope (Nikon eclipse T/2-CSU-X1) at 37□°C and 5% CO□. For lamellipodia dynamics, DIC time-lapse imaging were acquired every 2 sec for 10 min with a 63 X objective on a Zeiss Axio Observer Z.1 microscope.

### Migration image analysis

The Particle Image Velocimetry (PIV) technique was carried out utilizing the MatPIV software within the MATLAB environment (reference). For this examination, interrogation windows measuring 64 x 64 pixels (40.96 µm) and 32 x 32 pixels (20.48 µm) were employed, featuring a 50 % overlap. Vectors identified as outliers were manually excluded, and a local standard deviation filter was applied. The order parameter is described as the cosine of the angle between the velocity vector and the average direction of migration **(Figure S2E)**. This parameter ranges from 1, where the velocity vectors are aligned with the monolayer migration direction, to −1, for vectors directed opposite to the monolayer migration paths, and through 0, for vectors perpendicular to the monolayer migration. Profiles of the time-averaged velocity and order parameter were derived by processing the data in relation to the distance from the leading edge of migration, utilizing custom-written MATLAB scripts.

Employing in-house MATLAB script, spatial velocity correlation coefficients **(Figure S2E)** were determined by subtracting the average velocity from each frame prior to performing the autocorrelation analysis. The correlation coefficients were computed using these formulas:

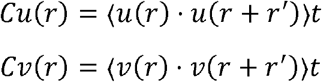

Where *C_u_* and *C_v_* represent to spatial correlation of the velocity components after detrending (removing the average velocity to highlight local velocity fluctuations) in the directions perpendicular and parallel to the migration direction. Here, u and v denote the detrended velocity fields, and **r** signifies a particular point where these fields (either u or v) are measured. The r + ѓ denotes a shifted position vector, where ѓ being the displacement vector from the reference point r. This displacement ѓ is correlated with the field at the position r. The notation for averages,(.)t, signifies the time average across each spatial point in the velocity field. To derive 1D function, these spatial velocity correlation functions were azimuthally averaged, therefore, *C_u_* and *C_v_* depend on ||r|| ( the norm of r). The correlation function was subsequently averaged over time for each migration sequence.

Protrusive structures, referred to as ‘fingers’, situated at the leading edges of the migrating cells, were manually identified through phase contrast video microscopy. Leader cells were identified as those positioned at the tip of these protrusions, characterized by their polarization, conspicuous lamellipodia, and crescent-shaped morphology. The quantification of the leader cells was performed manually and normalized against the length of the leading edge.

### Computation of Local velocity, cell orientation order parameter, directional persistence

The alignment of local velocities within fingers was assessed by calculating cos(θ), where θ ∈ [-180°, 180°] represents the angle between the leader cell mean velocity vector and the followers velocity vectors. Angular distributions were analyzed over 2-hour periods at 10-minute intervals. Velocity alignment was visualized using polar plots. To ascertain the local cell orientation order parameter, manual segmentation of the cells was performed, and the OrientationJ plugin in ImageJ was employed to discern the orientation of the cell’s major axis within the fingers. A custom MATLAB program was utilized to compute the angles between the major axis of the cells and the principal axis of these projections. The angular distributions were subsequently plotted concerning finger alignment. To determine directional persistence of cells, their migration trajectories were tracked manually using MtrackJ in ImageJ in phase contrast videos. Directional persistence was calculated as: Directional persistence = Cell displacement / Cell accumulated distance.

### Computation of distribution of cytoskeletal filaments, focal adhesions

To ascertain the distribution of cytoskeletal filaments within the monolayer and at the leader cell, fluorescence images were processed with a mask applying a threshold limit value set to 4. Custom-designed MATLAB scripts were utilized to quantify the average fluorescence intensity and standard deviation from the leading edge of the monolayer and the front edge of the leader cell across the image region delineated by the mask. To quantify the size and distribution of focal adhesions in the leader cells. Focal adhesions within the leader cells were masked applying a threshold limit value set to 4. Using ImageJ, size and number were quantified. Using home written MATLAB script, the leader cell region, extending from the front to the rear edge, was divided into five distinct sections, whereby the density of focal adhesions was quantified for each respective compartment.

### Tractions force Microscopy

Soft PDMS substrates were prepared using a 1:1 mixture of CyA:CyB (Dow Corning), spin-coated on a glass-bottom Petri dish at 500 rpm for 1 minute and cured at 80°C for 2 hours (Young’s modulus ≈ 15 kPa). The surface was silanized with 5% aminopropyl-triethoxysilane (APTES) in ethanol, rinsed, and dried. Red fluorescent carboxylated beads (200 nm) were applied and the substrates were coated with 50 µg/ml fibronectin. After overnight incubation, the culture insert was removed, and the dynamics of the cells and beads was captured by time lapse at a frequency of one frame per 10 minutes for 24 hours using time lapse video microscopy (Nikon Biostation). After 24 hours, cells were treated with sodium dodecyl sulfate (SDS). The bead displacement fields were determined via PIV (MATLAB scripts) and traction forces were calculated using Fourier transform traction cytometry (FTTC) in Fiji. Using home written MATLAB scripts traction forces were analyzed as described previously (Balasubramaniam et al., 2021), with heat maps and distance maps generated for force components.

### Computation of lamellipodia dynamics

To ascertain directional persistence, the spatial coordinates corresponding to the maximum extensions of the lamellipodia were manually identified over time, and the directional autocorrelation was evaluated utilizing a custom-developed MATLAB script. The angular deviations of the maximal lamellipodial extensions were calculated relative to the orientation of the leader cell over time. In order to ascertain the protrusion and retraction velocities of the lamellipodia, kymographs were produced from leader cells. These velocities were analyzed by measuring the slopes of the kymograph lines: Speed = Δx (μm) / Δt, where Δx (μm) represents the lengths of protrusion and retraction, and Δt denotes time. Persistence was further quantified as the duration over which protrusion and retraction persist. Additionally, the frequency of lamellipodia dynamics was determined by quantifying the number of protrusion and retraction events within a given time unit. To determine directional autocorrelation, time-lapse images were acquired every 10 min for 2 h. For each leader cell, the centroid trajectory was used to define the migration axis, corresponding to the major axis of movement. At each timepoint, lamellipodial protrusion coordinates were identified manually and compared with the major axis to obtain protrusion direction. Angles were extracted in MATLAB using *atan2* with angle unwrapping. Directional autocorrelation was then computed from the resulting angular time series using a custom-developed MATLAB script, allowing quantification of the temporal persistence of lamellipodial protrusion orientation.

### Lamellipodia and cryptic lamellipodia analysis

Cells were segmented manually using Cellpose. Cell contours were then extracted from the segmentation using custom-made MATLAB code, plotted with an increment of 5, and colour-codded according to time. They were then superposed over the first image of the corresponding movie. Cell contour dynamics kymographs were made using the methods and Matlab code (Marshall-Burghardt et al., 2024) adapted to our data. The original code is available on GitHub [https://github.com/HayerLab/Marshall-Burghardt-SciAdv2024/tree/main]. The contour was defined from the segmented masks and then divided in 180 sectors, which were tracked over time. The velocity of each sector relative to the direction normal to the local cell edge was computed to make the protrusion-retraction maps. Then, the average PBD-YFP/iRFP signal was measured in each sector, up to a depth of 20 pixels within the cell bulk. An external layer of 3 pixels was removed from the analysis to prevent strong signals from neighboring cells to interfere with the measurement. The PBD signal was then normalized as follows: the median of PBD signal was measured over the whole cell area and a sliding time window of 30 minutes duration. Then the PBD intensity values along the contour were divided by this median value. The edge velocity maps were smoothed both in space and time, using a disk mean filter of 2*2 (contour sectors*time frames) radius. The PBD maps were smoothed only in space using a mean filter of 5 contour sectors width.

To measure typical “contour fraction length” and time scale of protrusive areas **(Fig. Supp. S7)**, PBD kymographs were thresholded with a 1.1 threshold. The size of binarized areas in the kymographs was measured either along the horizontal direction at a given cell contour position or along the vertical direction at a given time frame, then all those sizes were pooled for a given cell and the average value of this pool was measured. We checked that qualitatively the trend – longer-lived and larger high-PBD sectors in WT than in KO cells was robust to a broad range of analysis parameters, namely: PBD threshold from 0.9 to 1.5; contour thickness (from the cell edge towards its interior) of 3 to 25 pixels; reference median PBD intensity value taken either over the whole cell surface or along the measured contour only.

### Statistical analysis

The data are depicted as the mean in dot plots or as mean ± SD in histograms, with at least three separate experiments conducted. Statistical analysis employed both the student’s t-test and the Mann-Whitney U test to address any non-normal distributions. These analyses were conducted using Python and GraphPad Prism. A p-value of less than 0.05 was marked as significant with one asterisk (*), while p < 0.01 and p < 0.001 were represented by two (**) and three (***) asterisks, respectively.

## Author Contributions

Conceptualization: RMM, BL, and SKP; methodology: BL, RMM, SV, SKP, SBM, TD, MA, RC, CT; software: LA, JA, JE, SKP; validation: BL, RMM; formal analysis: SKP, SBM; Investigation: SKP, CT; resources: BL, RMM, SBP; data curation: RMM, SKP; writing-original draft: RMM, SKP; writing-review and editing: RMM, BL, SKP, SBP, JA, LA; visualization RMM, SKP, SBP, CT; supervision: BL, RMM; project administration: BL, RMM; and funding BL, RMM, SV.

## Acknowledgements

This work was supported by the European Research Council (Grant No. Adv-101019835 to BL), LABEX Who Am I? (ANR-11-LABX-0071 to BL and RMM), the Ligue Contre le Cancer (Equipe labellisée 2019 to RMM), the Alexander von Humboldt Foundation (Alexander von Humboldt Professorship to BL), the Agence Nationale de la Recherche (“STRATEPI” DFG-ANR-22-CE92-0048 to RMM) and (ANR-20-CE13-0024-01, ANR-21-CE13-0018-01 to SV). JE was supported by the Australian Research Council (FL230100100 and DP220103951) and the Deutche Forschungsgemeinschaft (DFG, German Research Foundation)-553948485.

We would like to thank all the members of the “Cell adhesion and Mechanics” team for helpful discussions. We thank Gautham Hari Narayana for initial help, training and generation of Vimentin KO cell lines, Nicolas Borghi and Cecile Leduc for all the helpful discussions, as well as Xavier Baudin and Nicolas Moisan for their help with imaging. We acknowledge the ImagoSeine core facility of the IJM, a member of IBiSA and France-BioImaging (ANR-10-INBS-04) infrastructures.

