## Supplemental Figures for "Vimentin bridges scales to convert polarized cell locomotion into coordinated collective migration"

### Multiple length scales effects of vimentin on collective cell migration

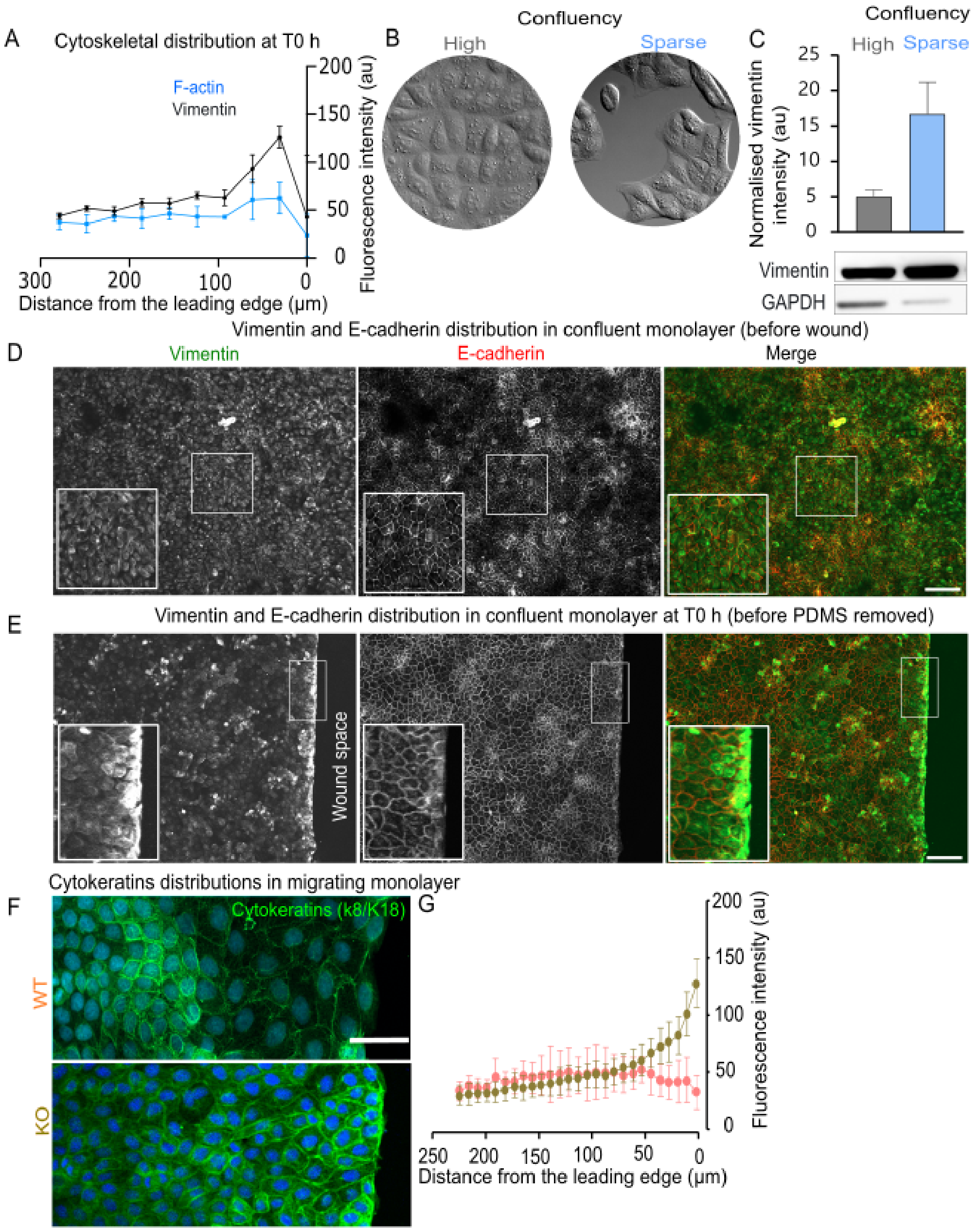


#### Figure S1: Distribution of cytoskeletons in migrating monolayers

**(A)** Quantification of the distribution of vimentin and F-actin stainings at t 0 h after removal of physical constraint. Sample sizes n = 3 monolayers. **(B, C)** Vimentin protein levels in confluent and sparse culture as analyzed by Western blotting, histograms indicate means ± SD. The histogram represents the quantification of vimentin levels normalized to GAPDH. **(D)** Immunostainings of vimentin and E-cadherin in confluent monolayers without free edge. Scale bars, 100 µm. **(E)** Immunostainings of vimentin and E-cadherin distribution before removal of physical constraint. Scale bars, 100 µm. **(F, G)** Cytokeratin stainings in WT and KO monolayers at t 24h. The dots represent average values binned every 15 µm and bars indicate SDs. Sample sizes n=10 monolayers for WT and KO monolayers. Scale bars, 50 µm. Sample sizes were representative of three independent experiments.

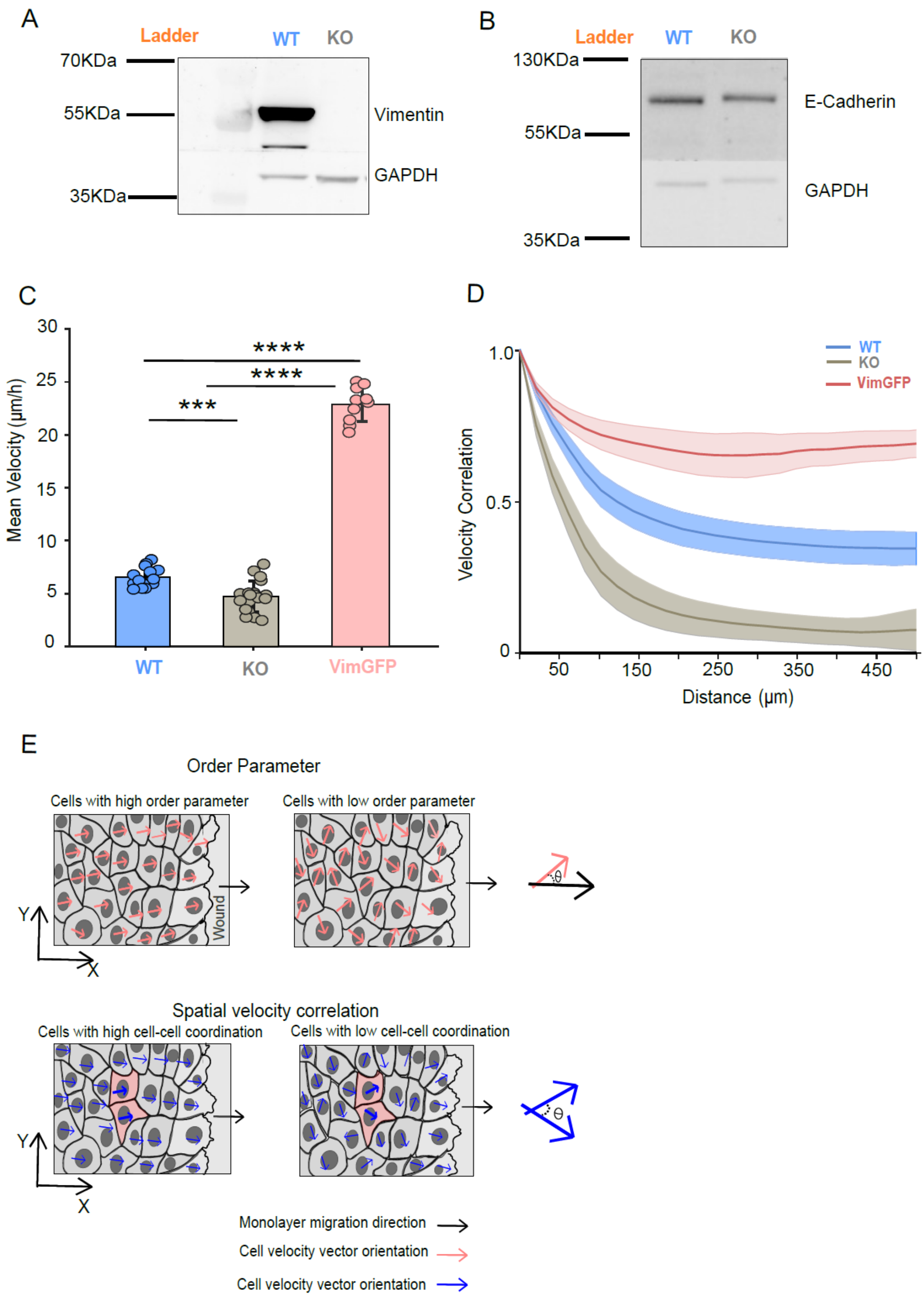


**Figure S2: Vimentin does not affect E-cadherin expression.**

**(A, B)** Western blot analysis of protein extracted from WT and KO cells showing the total depletion in vimentin in KO cells and unchanged E-cadherin levels compared to WT conditions.

**(C)** Mean velocity of WT, KO and Vim-GFP cells in the migrating monolayer over 15 hours. Sample sizes n=11 (WT), n-19 (KO), n=15 (Vim-GFP). N= 3 independent experiments (WT and KO). N=2 Independent experiments (Vim-GFP). The data points indicate averaged values, and bars represent the mean ± SD, Mann-Whitney comparisons. **(D)** Average spatial velocity correlation function over distance for WT, KO and Vim-GFP monolayers. Thick lines and shaded regions represent mean and standard deviation, respectively. Sample sizes n=11 (WT), n-19 (KO), n=15 (Vim-GFP). N= 3 independent experiments (WT and KO). N=2 Independent experiments (Vim-GFP). **(E)** Velocity order parameter analysis. The cartoon illustrates the velocity order parameter, calculated as cos($\theta$), where $\theta$ is the angle between each local cell velocity vector (Orange vector) and the average monolayer migration direction (Black vector). In directionally coordinated monolayers, local velocity vectors remain aligned with the average monolayer migration direction, producing a high order parameter. In poorly coordinated monolayers, local velocity vectors deviate from the migration direction, producing a lower order parameter. **(F)** Spatial velocity correlation analysis. The spatial velocity correlation function quantifies how local velocity magnitudes (blue arrows) correlate between pairs of positions (cells in orange) separated by a distance 'r'. In coordinated monolayers, velocity magnitudes remain correlated over longer distances, producing a slow decay of the correlation function. In poorly coordinated monolayers, velocity magnitudes decorrelate over shorter distances, producing a faster decay.

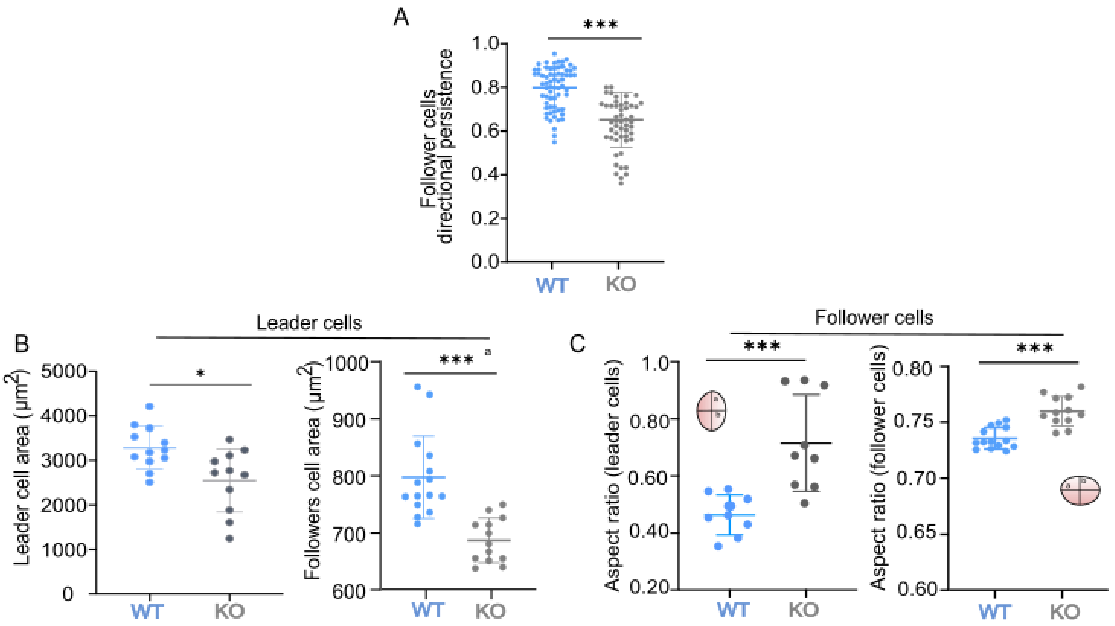


**Figure S3: Vimentin maintains cells morphology and involved in directed migration**

**(A)** Directional persistence of followers. Each color filled dots represents a follower cell. Colored bars indicate average value and SDs. Mann-Whitney comparison, n=66 followers (WT) and n = 52 (KO). **(B)** Quantification of leader cell and follower cell area. Each data point representative of individual leader and follower cells. Bars represent the mean values ± SDs. Mann-Whitney comparisons, cell area n=12 leaders (WT) and n=11 leaders (KO). **(C)** Quantification of leader cells and follower cells aspect ratio (major axis (a) /minor axis (b)). Each data point represents an individual leader and follower cells. Bars represent the mean values ± SDs. Mann-Whitney comparisons, n=16 leaders (WT) and n= 16 leaders (KO). Sample sizes were representative of three independent experiments.

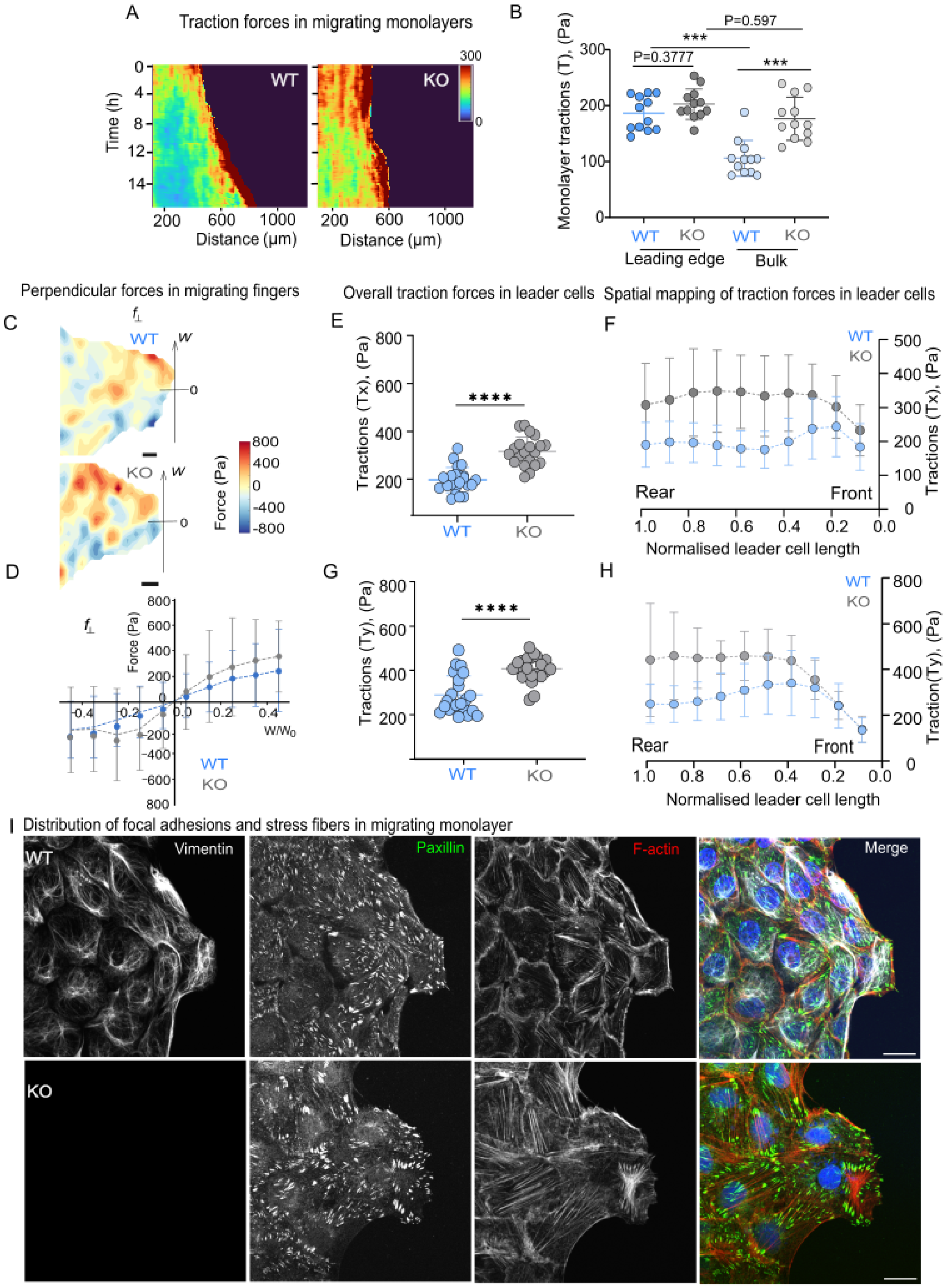


**Figure S4: Vimentin controls traction forces and distribution of FAs and stress fibers in the migrating monolayers**

**(A)** Kymographs show migrating monolayers with traction forces. Red indicates strong tractions; blue represents lower tractions. **(B)** Average traction forces at the leading edge (100µm) and within the bulk (200µm) of the migrating monolayer. Each data point represents a monolayer. Bars indicate the mean values ± SDs. Mann-Whitney comparisons, n= 12 monolayers (WT), n= 12 monolayers (KO). **(C)** Heat maps displaying the distribution of perpendicular forces in WT and KO fingers, along with the corresponding forces normalized to finger width. Mann-Whitney comparisons, n= 12 monolayers (WT), n = 12 monolayers (KO). **(E, G)** Quantification of traction forces over a period of 2 hours in whole leader cells in both Tx and Ty directions. Each data point indicates the leader cell. Bars indicate the mean values ± SDs. Mann-Whitney comparisons n= 21 leaders (WT), n= 20 leaders (KO). **(F, H)** Quantification of distribution of traction forces over a period of 2 hours in leader cells in both Tx and Ty directions. Each data point indicates average tractions binned 0.1 units. Bars and broken lines indicate average values (trend) ± SDs. Mann-Whitney comparisons n= 21 leaders (WT), n= 20 leaders (KO). **(I)** Immunostainings of vimentin (gray), Paxillin (green), F-actin (red) in migrating monolayers at t 24 hrs. Scale bars 50µm. Sample sizes were representative of three independent experiments.

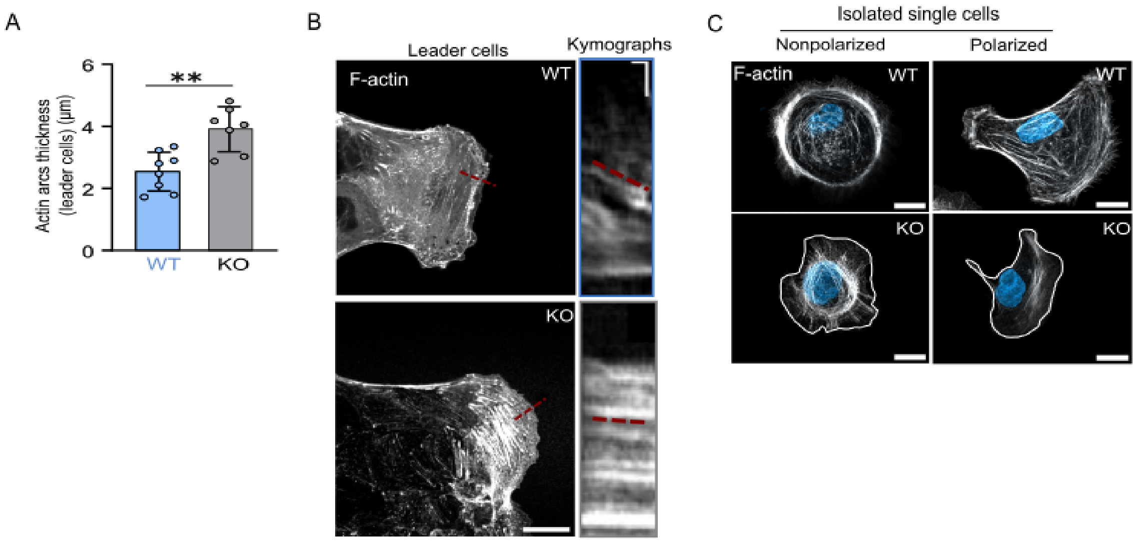


**Figure S5: Vimentin required for maintaining actin arc structure and its dynamics.**

**(A)** Quantification of actin arcs thickness in leader cells. The bars represent mean values ± SDs, Mann-Whitney comparisons, n=8 (WT), 7 (KO) leader cells. **(B)** Images and corresponding kymographs of supplementary video5, showing actin arc dynamics-in WT and in KO leader cells. The dashed red line indicates retrograde F-actin speed of corresponding structures. Scale bars, 20 µm (main images) and 30 s/2 µm (*x*/*y* axes) in the kymographs. **(C)** F-actin staining in isolated cells after 3 hours of seeding depicting the structure of F-actin in individual non-polarized and polarized cells. Scale bars, 20 µm. Sample sizes were representative of three independent experiments.

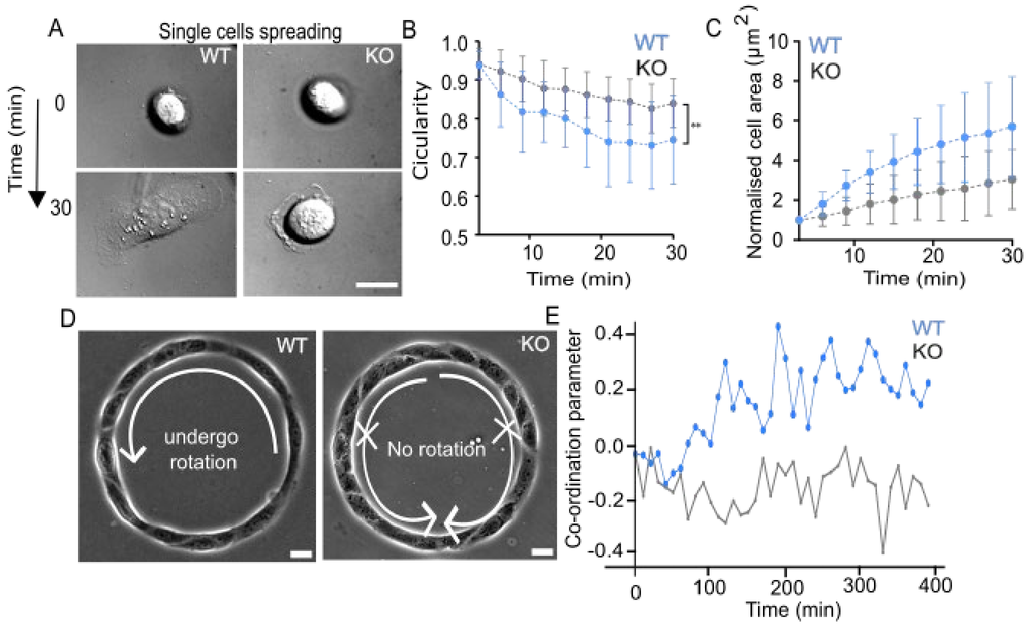


**Figure S6: Vimentin required for maintaining the front-rear polarity in epithelial cells.**

**(A)** Images of single cells spreading on conditioned substrates corresponding to supplementary video8. Scale bars, 10μm. **(B,C)** Quantification of the evolution of spreading cell shape and cell area over time. The dotted line and bars indicate the mean values ± SDs. Mann-Whitney comparisons, n=12 cells (WT), and n=14 cells (KO). **(D, E)** Phase contrast images of a train of cells in WT and KO single rings. The white arrow marks represent direction of rotation. The graph indicates the mean values of coordination parameters in WT and KO. Sample sizes n= 10 single rings (WT) and n=10 single rings (KO). Samples sizes C, F and D were representative of three and two independent experiments.

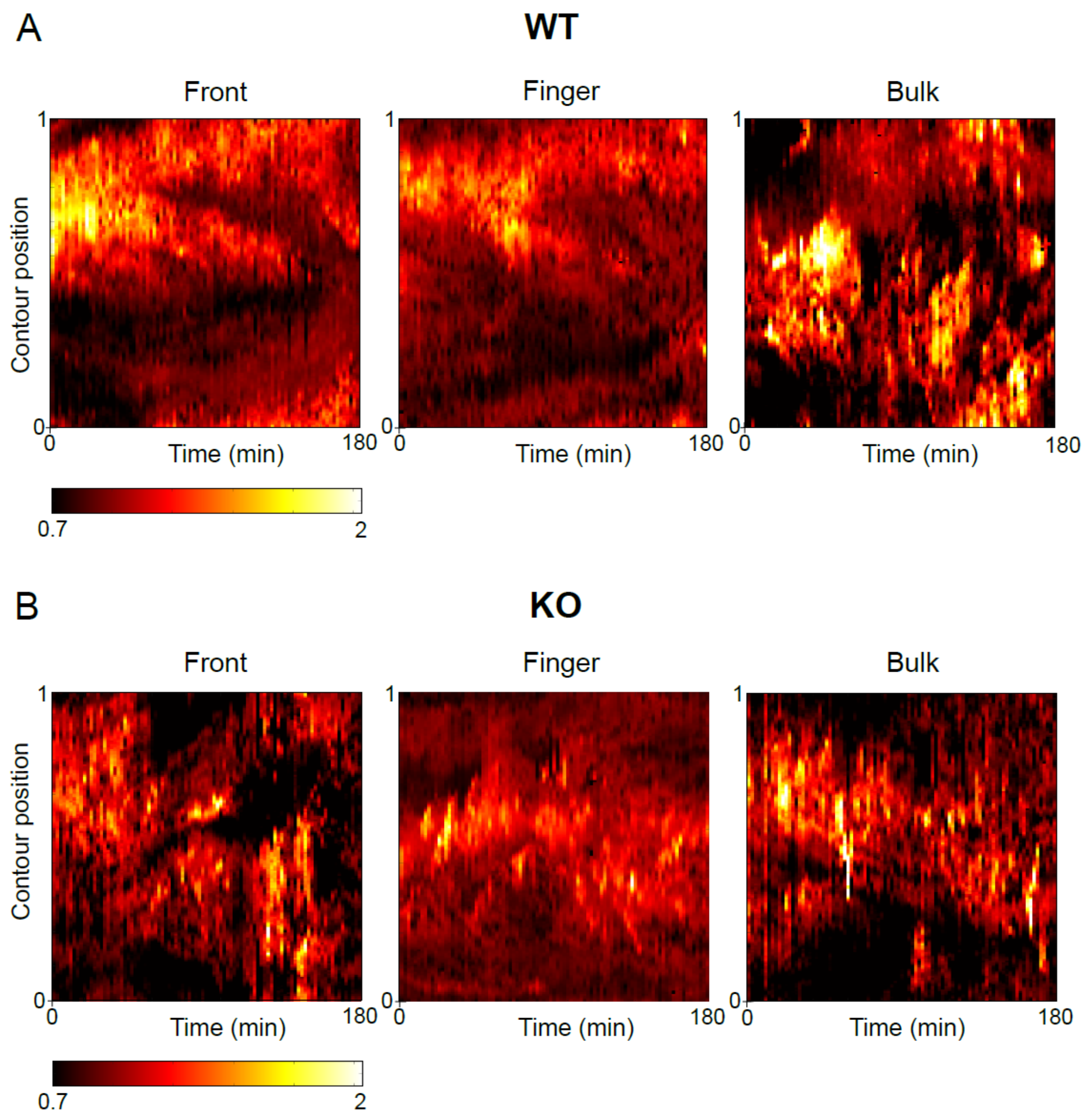


**Figure S7: Evolution of the PBD-YFP signal at the periphery of front, finger and bulk cells.**

Maps of PBD-YFP signal 20 pixels from the edge around the linearised cell for front, finger and bulk WT**(A)** and Vim KO cells **(B)** shown in Figure 7.
